# Glia-enriched stem-cell 3D model of the human brain mimics the glial-immune neurodegenerative phenotypes of multiple sclerosis

**DOI:** 10.1101/2024.06.20.597748

**Authors:** Francesca Fagiani, Edoardo Pedrini, Stefano Taverna, Elena Brambilla, Valentina Murtaj, Paola Podini, Francesca Ruffini, Erica Butti, Clarissa Braccia, Annapaola Andolfo, Roberta Magliozzi, Lena Smirnova, Tanja Kuhlmann, Angelo Quattrini, Peter A. Calabresi, Daniel S. Reich, Gianvito Martino, Paola Panina-Bordignon, Martina Absinta

## Abstract

The role of central nervous system (CNS) glia in sustaining self-autonomous inflammation and driving clinical progression in multiple sclerosis (MS) is gaining scientific interest. We applied a single transcription factor (*SOX10*)-based protocol to accelerate oligodendrocyte differentiation from hiPSC-derived neural precursor cells, generating self-organizing forebrain organoids. These organoids include neurons, astrocytes, oligodendroglia, and hiPSC-derived microglia to achieve immunocompetence. Over 8 weeks, organoids reproducibly generated mature CNS cell types, exhibiting single-cell transcriptional profiles similar to the adult human brain. Exposed to inflamed cerebrospinal fluid (CSF) from MS patients, organoids properly mimic macroglia-microglia neurodegenerative phenotypes and intercellular communication seen in chronic active MS. Oligodendrocyte vulnerability emerged by day 6 post-MS-CSF exposure, with nearly 50% reduction. Temporally-resolved organoid data support and expand on the role of soluble CSF mediators in sustaining downstream events leading to oligodendrocyte death and inflammatory neurodegeneration. Such findings support implementing this organoid model for drug screening to halt inflammatory neurodegeneration.

## INTRODUCTION

Multiple sclerosis (MS) is the most common chronic inflammatory, demyelinating, and neurodegenerative disease of the central nervous system (CNS), extensively involving the brain, spinal cord, and optic nerve.^1^ Although currently approved disease-modifying treatments are effective in modulating peripheral immunity and reducing relapse-associated inflammatory activity, they fail to prevent disease progression and disability in a large proportion of patients.^2^ To halt MS clinical progression, the role of the macroglia-microglia axis in sustaining chronic inflammation (e.g., at the leading edge of chronic active lesions)^3^ and in limiting the restoration of tissue homeostasis is increasingly attracting scientific and pharmacological interest.^2^

In the absence of preclinical models of smoldering MS lesion formation and maintenance, we here report a 3-dimensional (3D) stem cell-derived model of the human brain as a reproducible and valuable tool to experimentally probe human CNS macroglia-microglia crosstalk. To be useful for this purpose, we envisioned that such a stem-cell based model would need to satisfy the following requirements: (1) harboring cellular lineage diversity to be as much as possible representative of the human brain; (2) displaying a high proportion of mature cells; and (3) achieving single-cell transcriptional similarity with neurodegenerative glial phenotypes seen in MS lesions.^3^

Our cellular platform consists of 3D multilineage submillimetric organoids integrating neural progenitor cell (NPC)-derived self-organizing forebrain organoids with human induced pluripotent stem cell (hiPSC)-derived microglia (also termed as assembloids;^4^ **Figure 1A**). They contain a mosaic of intermingled CNS cells (neurons, astrocytes, oligodendrocyte lineage cells, and microglia) that spatially and temporally overlap, allowing functional crosstalk among the different cell types. Recognizing major limits in the generation of long-lasting 2D cultures of oligodendrocytes (OLs), here, to accelerate oligodendrogenesis from hiPSC-derived NPC, we applied a single transcription factor (*SOX10*) based-protocol that promotes the commitment and the generation of a heterogenous and stable pool of oligolineage cells, including oligodendrocyte progenitor cells (OPCs) and myelin basic protein-positive (MBP^+^) OLs, with a global gene-expression profile comparable to primary human OLs.

**Figure 1.**
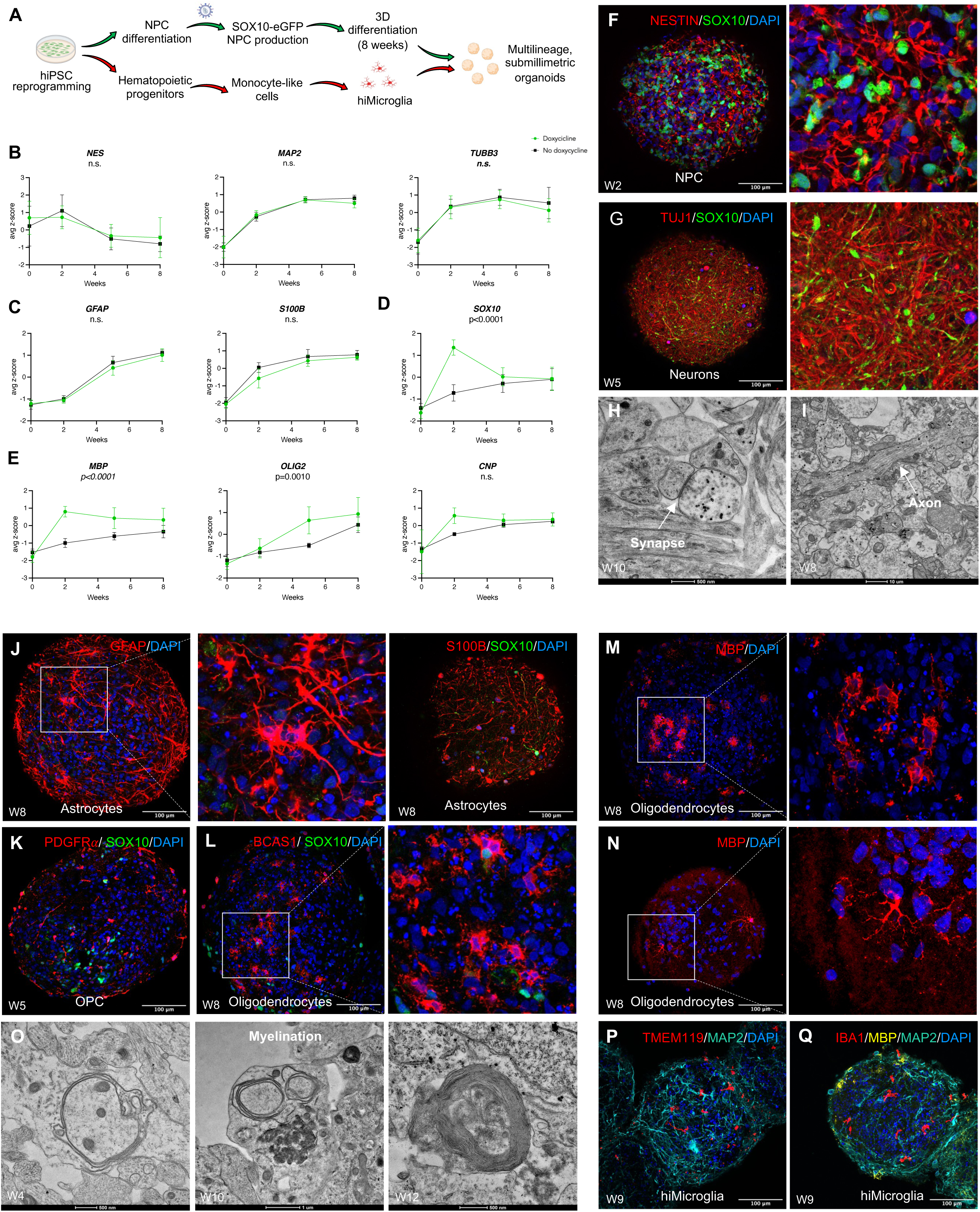
SOX10 transcription factor accelerates oligodendrogenesis in hiPSC-derived organoids. (A) Schematic representation of the protocol for generating submillimetric organoids and integrating hiPSC-derived microglia. (B-E) Gene expression (normalized to *GAPDH*) of *NES*, *TUBB3*, *MAP2, GFAP, S100B, SOX10*, *MBP*, *OLIG2*, and *CNP* in SOX10-eGFP organoids — exposed and not exposed to doxycycline for inducing *SOX10* expression — at 2, 5, and 8 weeks of differentiation, as determined by RT-qPCR (mixed effect analysis, followed by multiple comparison test) (for doxycycline-treated organoids: RNA samples each consisting of ∼150 organoids from 5 hiPSC lines from 2 differentiation experiments; for doxycycline-untreated organoids: RNA samples each consisting of ∼150 organoids from 5 hiPSC lines from 1 differentiation experiment). Data are represented as average z-score ± SD. (F) Immunostaining of Nestin^+^ and SOX10^+^ cells in doxycycline-treated organoids after 2 weeks of differentiation (magnification, 30X). (G) Immunostaining of Tuj1^+^ and SOX10^+^ cells in doxycycline-treated organoids after 5 weeks of differentiation (magnification, 30X). (H) Electron microscopy images of doxycycline-treated organoids after 10 weeks of differentiation showing a synaptic junction containing vesicles (white arrowhead). (I) Electron microscopy images of doxycycline-treated organoids after 8 weeks of differentiation showing axonal projections (white arrowhead). (J) Immunostaining of GFAP^+^ (30X) and S100B^+^ cells (30X) in cryosections from 8-week-old doxycycline-treated organoids. (K) Immunostaining of PDGFRα^+^ and SOX10^+^ cells in cryosections from 5-week-old doxycycline-treated organoids (30X). (L) Immunostaining of BCAS1^+^ and SOX10^+^ cells in cryosections from 8-week-old doxycycline-treated organoids (30X). (M-N) Immunostaining of MBP^+^ cells in cryosections from 8 weeks-old doxycycline-treated organoids (30X). (O) Electron microscopy images of organoids showing myelination at different time points (additional examples in **Figure S2C**). (P) Immunolabeling of TMEM119^+^ cells and MAP2^+^ in cryosections from 9-week-old organoids (30X). (Q) Immunolabeling of IBA1^+^ cells, MBP^+^, and MAP2^+^ cells in cryosections from 9-week-old organoids (30X). *Abbreviations:* NPC: neural precursor cells.

Compared to other cellular models, the major advantage of this 3D platform relies on the presence of mature glial populations, including hiPSC-derived microglia, and neurons capable to respond to complex inflammatory stimuli and partially resembling the inflammatory environment of the human MS brain. The presence of all these glial subtypes, including the immune counterpart, is of key relevance to dissect clinically relevant mechanisms driving the unresolved inflammation and propagating tissue injury of chronic active MS lesions^3,5^ and to provide a drug-discovery platform for high-throughput screens of compounds tackling inflammatory demyelination and neurodegeneration.

## RESULTS

### Ectopic expression of SOX10 transcription factor promotes oligodendrogenesis in hiPSC-derived organoids

To foster the generation of human OLs, we generated a lentiviral vector encoding the SOX10 coding sequence and the green fluorescent protein (GFP) as a reporter gene under the control of a doxycycline-inducible promoter (**Figure S1A**). Human iPSC-derived forebrain-patterned NPCs (n=5, 2 from non-neurological controls and 3 from individuals with MS, **Table S1**) were infected with SOX10-eGFP-expressing lentivirus, and the stably infected cells were sorted by fluorescence-activated cell sorting (FACS)-gating on GFP^+^ cells (**Figure S1B**). To recapitulate the complex cellular environment of the human brain *in vitro,* we sought to generate controlled-sized, submillimetric (∼450 μm-diameter), forebrain organoids (**Figure S1C and S1D**), consisting of neurons, astrocytes, and OLs, by optimizing previously published protocols.^6^ We derived organoids from SOX10-eGFP NPCs exposed to glial induction medium (GIM), supplemented with triiodo-L-thyronine (T3), recombinant human neurotrophin-3 (rhNT3), recombinant human PDGF BB homodimer (rhPDGF), smoothened agonist (SAG), 2’-o-dibutyryladenosine 3’, 5’-cyclic adenosine monophosphate (dbcAMP) analog, insulin-like growth factor-1 (IGF-1), ascorbic acid (AA), and 1 μg/mL doxycycline from day 1 to day 3 (as schematized in **Figure S1E**) (see **STAR METHODS** for a detailed description).^7,8^ On day 5, medium was replaced with glial differentiation medium (GDM), supplemented with T3, rhNT3, IGF-1, AA, and dbcAMP, to promote OPC proliferation and maturation. Organoids were cultured in GDM until the end of the differentiation protocol. Doxycycline at the final concentration of 1 μg/mL was added to GDM medium from day 5 until day 14 (**Figure S1E**).^7,8^ On day 14, 10 ng/mL BDNF and GDNF were added to support neuronal and glial trophism. Selected organoids from each cell line were not exposed to doxycycline (therefore no *SOX10* induction) and considered as negative controls.

First, to assess the maturation stages of organoids during the 8-week differentiation protocol, NPCs as well as 2-, 5-, and 8-week-old organoids (from 5 hiPSC lines) were collected for RNA extraction. The expression of a panel of neuronal, astrocyte, and oligodendrocyte markers was analyzed by real-time quantitative polymerase chain reaction (RT-qPCR); their longitudinal expression was compared in organoids exposed *vs* not exposed (negative control) to doxycycline for 14 days. The gene expression of *NES*, a neural stem/progenitor cell marker (*NESTIN*), remained similar to NPCs until 2 weeks of differentiation when the expression started to decrease (**Figure 1B**). At the end of the differentiation protocol, the remaining *NES* expression indicates the presence of a small population of NPCs, as well as other proliferating cell types, such as OPCs and astrocytes. Accordingly, immunofluorescence analyses showed a net reduction in Nestin^+^ and Pax6^+^ cells through the differentiation process (**Figure S1I** and **S1J**). Compared to NPCs, after 2 weeks of differentiation, we also observed a significant increase in the gene expression of both neuronal (*i.e., MAP2*, *TUBB3*) (**Figure 1B**) and astrocyte markers (*i.e., GFAP, S100B*) (**Figure 1C**). As expected, no differences in the expression of neuronal and astrocyte markers were observed between organoids stimulated with doxycycline compared to the non-stimulated ones. To confirm the doxycycline-induced increase in *SOX10* expression, we analyzed *SOX10* mRNA expression, demonstrating a significant increase after 2 weeks of differentiation in organoids exposed to doxycycline compared to negative control conditions (**Figure 1D**). As expected, after doxycycline discontinuation, *SOX10* expression started to decrease. With respect to OPC and mature oligodendrocyte lineage markers, a significant increase in *MBP* and *OLIG2* gene expression was observed in organoids exposed to doxycycline compared to untreated organoids (**Figure 1E**). We observed significantly higher *MBP* mRNA levels after 2 weeks of differentiation in organoids stimulated with doxycycline, suggesting that SOX10 induction directly promoted its expression. As expected, *OLIG2* gene expression started to increase after 5 weeks of differentiation. Such evidence is consistent with data indicating that *OLIG2,* besides acting as an upstream regulator of *SOX10* expression in OPCs, also directs chromatin remodeling to potentiate the expression of several my-elin-related factors that initiate OPC differentiation into OLs (**Figure 1E**).^9,10^

Next, by immunofluorescence analysis, we observed the maturation of NPCs (**Figure 1F**) into an intricate network of Tuj1^+^ (**Figure 1G**) and MAP2^+^ (**Figure S1G and S1H**) neurons after 5 weeks of differentiation (without any cortical layer-like pattern), with axonal projections (**Figure 1I and S2B**) evident on transmission electron microscopy. Ultrastructural analysis revealed the presence of synaptic junctions, capturing processes of synaptic vesicle fusion with docked vesicles in contact with the plasma membrane in the active zone (**Figure 1H and Figure S2A**). To confirm functional interactions between neurons in organoids, we performed electrophysiological recordings on 12-week-old organoids using whole-cell patch clamp recordings. Repetitive firing of action potentials (indicating a mature, neuronal-like functional profile) was observed in response to suprathreshold current injection in 10 out of the 49 cells tested (20.4%; **Figure S1F**) (mean spike frequency was 10.8±1.2 Hz). These neurons displayed sizeable Na^+^ and K^+^ currents in response to step voltage commands from -70 mV to -10 mV (**Figure S1F**), with average peak amplitudes of 2864±470 pA and 1560±338 pA, respectively. In 7 of the neurons, we also recorded a mixture of spontaneous excitatory and inhibitory postsynaptic currents (**Figure S1F**). Astrocytes were also clearly identified in 8-week-old organoids, as showed by the presence of GFAP^+^ and S100B^+^ cells with typical astrocyte morphology (**Figure 1J**).

Concerning the OLs, by 5 weeks of differentiation, PDGFRa^+^ cells were detected, indicating OPC proliferation (**Figure 1K**). By 8 weeks of differentiation, we observed OL maturation indicated by the presence of breast carcinoma amplified sequence 1 (BCAS1) ^+^ cells, accounting for a population of newly formed, myelinating oligodendrocytes marking regions of active myelin formation (**Figure 1L**). In addition, we detected myelin basic protein (MBP)^+^ cells with a highly branched morphology after 8 weeks of differentiation (**Figure 1M and 1N**). As proof of myelination, SOX10-eGFP organoids treated with doxycycline were imaged with transmission electron microscopy at 4, 8 and 12 weeks of differentiation, demonstrating the presence of various stages of myelination, including early myelination processes and myelin wraps surrounding axons (**Figure 1O and S2C**).

Since microglia originate from yolk sac erythromyeloid progenitors generated during primitive hematopoiesis,^11^ to achieve an immunocompetent organotypic model, we generated in parallel iPSC-derived human microglia (hiMicroglia), according to the two-step protocol of Abud *et al.*^12^ On day 34 of differentiation, microglia morphology (small cell body and ramified processes), as well as the presence of typical human microglial markers (CD11B, CD45, and TMEM119) was confirmed by immunofluorescence in 2D culture of hiMicroglia (**Figure S1K and S1L**). Then, hiMicroglia were incorporated in 8-week-old organoids producing hiAssembloids.^4^ After 24h, microglia spontaneous migration and integration into the neuron-glial network was observed, as confirmed by immunostaining on 15 μm-thick cryosections showing IBA1^+^ and TMEM119^+^ cells intermingled with neurons and glia (**Figure 1P, 1Q, S1M, S1N, S1O, and S1P)**, thereby supporting the widely described microglia mobility and motility.

### Single-cell RNA sequencing unraveled the cellular diversity of organoids

To comprehensively characterize the cellular landscape of our model, we performed single cell RNA-sequencing of the 8-week-old SOX10-eGFP expressing organoids with hiMicroglia (both exposed and nonexposed to doxycycline) and organoids obtained based on a standard protocol (GIBCO, a well-established and commercially available protocol to produce 3D brain organoids). After standard preprocessing, we generated data for a total of 29,319 single cells from 9 individual samples, generated from 5 independent stem cell lines (**Table S2**). Overall, 22,089 distinct transcripts were detected. Unsupervised clustering identified 17 initial clusters, rendered as uniform manifold approximation and projection (UMAP) (**Figure 2A**). The annotation was performed using known lineage marker genes (**Figure 2B and S3A**). For uncertain assignment, we relied on the hallmark lineage genes of each cluster (top 100 positive differentially expressed genes against all the other cells) and performed over representation analysis (ORA), using a panel of annotations (KEGG 2021, Human MSigDB Hallmark 2020, Reactome 2016, Human Gene Atlas Azimuth Cell Types 2021).

**Figure 2.**
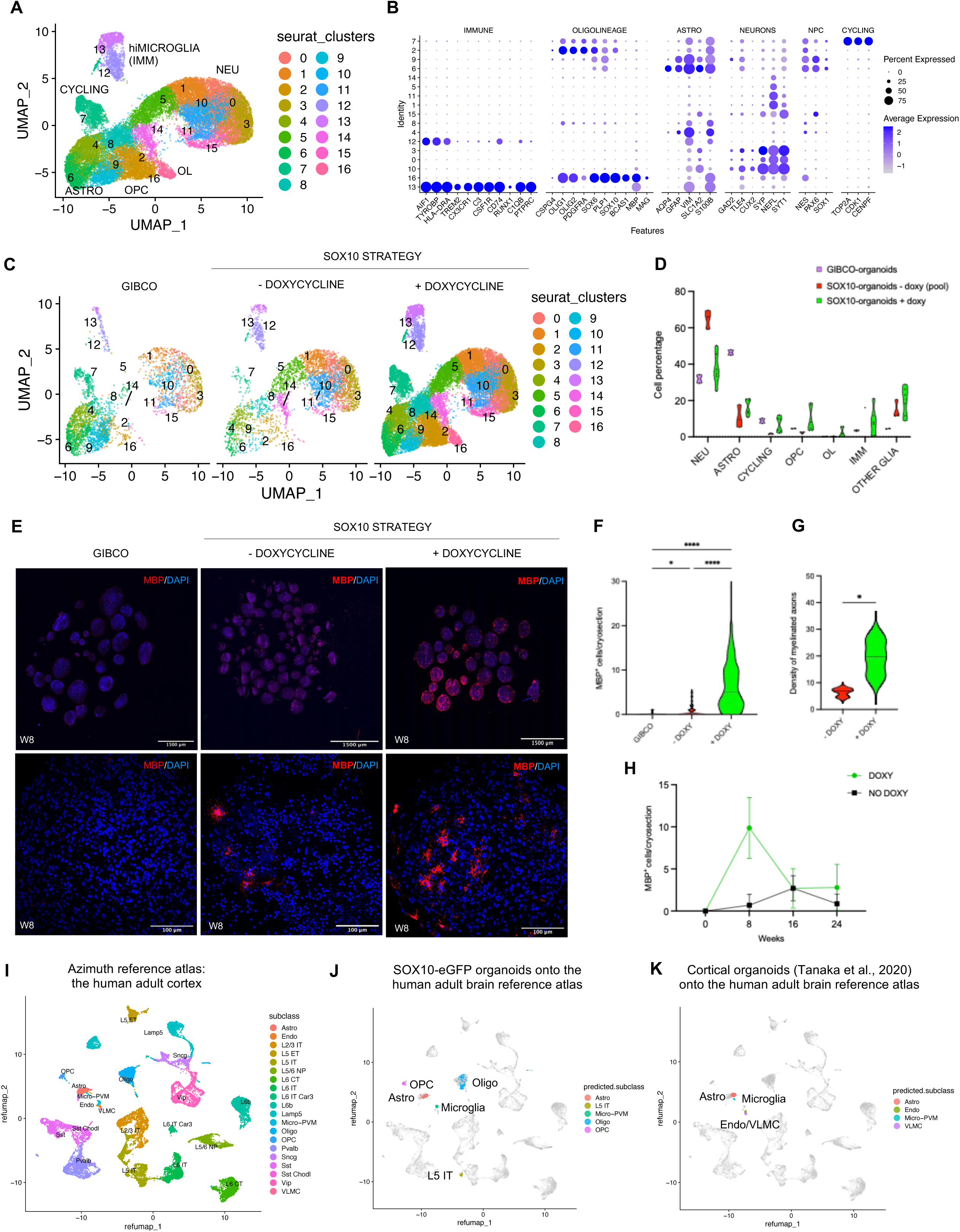
Single-cell RNA sequencing unraveled the cellular diversity of organoids, recapitulating the cell diversity of the human adult cortex. (A) scRNA-seq clustering of 29,319 cells, labelled based on known lineage markers and visualized as UMAP plot, from 8-week-old organoids deriving from SOX10-eGFP-expressing NPCs that were exposed to doxycycline for 2 weeks to promote SOX10 induction. Each dot corresponds to a single cell and each color to a cell cluster. (B) Dot plot depicting selected differentially expressed genes for each cluster and associated cluster labelling. Dot size corresponds to the percentage of cells expressing the gene in each cluster, and the color represents the average expression level. (C) scRNA-seq UMAP clustering from organoids produced based on the GIBCO protocol (*n*=3,660 cells) and SOX10 strategy, exposed (*n*=21,380 cells) and not exposed to doxycycline (*n*=4,279 cells), at 8 weeks of differentiation. (D) Violin plot showing the percentages of different cell populations by protocol. (E) Immunostaining of MBP^+^ cells in 15 µm-thick cryosections containing multiple 8-week-old organoids produced based on the commercially available GIBCO protocol *vs* our SOX10 strategy both exposed and not exposed to doxycycline (top row: 10X; bottom row (magnified view of one organoid): 30X). (F) Quantification of branched MBP^+^ cells in cryosections (estimated cryosection area=0.2 μm^2^) from 8-week-old organoids produced based on the different (*n*=6 samples, each consisting of 30 cryosections derived from 5 hiPSC lines). (G) Quantitative analysis of myelinated axons/1000 μm^2^ area for doxycycline-stimulated organoids *vs* doxy-minus organoids (*n*=3). (H) Quantification of branched MBP^+^ cells in cryosections from organoids at 8, 16, and 24 weeks of differentiation (30 cryosections for each time point from 1 hiPSC line). (I) Reference atlas of the adult human primary motor cortex from cases without neurological disease (Azimuth).^17^ (J) Mapping of 3D organoids single-cell data (color-coded) onto a reference atlas of the adult human primary motor cortex (gray dots, H).^17^ (K) Mapping of single-cell transcriptome profiles of human cortical organoids from eight different protocols collected from public resources (differentiation range ∼60/70 days) (dataset metanalysis available in Tanaka et al.^28^) on the reference atlas of the adult human primary motor cortex (gray dots, H).^17^ *Abbreviations:* NEU:neurons; ASTRO:astrocytes; OPC:oligodendrocyte precursor cells; OL:oligodendrocytes; NPC:neural precursor cells; hiMICROGLIA (IMM):myeloid immune cells (hiPSC-derived microglia); OTHER GLIA:immature astrocytes; + DOXY:doxycycline-treated; - DOXY:not treated with doxycycline; Micro-PVM:microglia-perivascular macrophages; Endo:endothelia; VLMC:vascular and leptomeningeal cells.

We identified distinct populations of neurons, astrocytes, OPCs, OLs, and immune cells. We observed the presence of different types of neurons, such as *GAD2^+^* inhibitory and excitatory neurons, some showing upper (*CUX2*^+^) *vs* lower (*TLE4^+^*) cortical layer specification (**Figure 2B**). In accordance with the forebrain-oriented differentiation pattern, we did not observe the presence of dopaminergic neurons. We also observed the presence of mature astrocytes expressing typical astrocyte lineage markers, such as *AQP4*, *GFAP*, *VIM,* and *S100B* (**Figure 2B**). One oligolineage cluster (cluster 2) was annotated as OPCs (*SOX10^+^*, *OLIG1^+^, OLIG2^+^, and PDGFRa^+^)* (**Figure 2B**). As expected, myelin genes (*i.e., MBP, PLP1,* and *MAG)* and *BCSA1* were differentially expressed in the oligolineage cluster (cluster 16) than in the OPC cluster (cluster 2) (**Figure 2B**). As expected, *SOX10^+^* expressing cells were mainly restricted to oligolineage cells and some of the cycling cells of cluster 7 (**Figure S3B)**. The hiMicroglia clusters (cluster 12 and 13) showed similar patterns of marker expression (*AIF1*, *TYROBP*, *HLA-DRA, TREM2, CX3CR1, C3, CSF1R, C1QB and PTPRC*) as the human microglia (**Figure 2B**). In addition, we observed the presence of a pool of cycling and proliferating cells with enriched expression of cell-cycle-related genes (*TOP2A*, *CDK1,* and *CENPF*) (**Figure 2B**). Cell-cycle scoring confirmed these cells as being in the G2M/S phase, whereas the other clusters were mainly in the G1 phase of the cell cycle (**Figure S3B**). The lists of the top 100 positive differentially expressed genes for each cluster against all the others are provided as supplementary information (**Table S3**).

In line with prior reports,^13–15^ compared to astrocytes derived from control hiPSC-lines, MS-derived astrocytes showed an enrichment in self-antigen presentation, extracellular matrix organization, inflammatory signaling (*i.e.,* interferon signaling, IL4 and IL13 signaling) as well as response to hyoxia (**Table S4**). Compared to control-derived neurons, MS-derived neurons showed an enrichment in cell energy-related terms (*i.e.,* mitochondrial electron transport, oxidative phosphorylation) and cholesterol biosynthesis (**Table S4**). When compared to control-derived OPC, MS-derived OPC showed an enrichment in cellular response to oxidative stress (**Table S4**). No major differences were seen in the transcriptional profile of MS vs control-derived oligodendrocytes.

To test the efficiency of SOX10 strategy in promoting OL maturation, we compared OL differentiation in 8-week-old organoids obtained based on GIBCO protocol with our SOX10-eGFP expressing organoids, stimulated *vs* not stimulated with doxycycline. The GIBCO protocol did not produce mature OLs (**Figure 2C**). On the other hand, organoids generated using our SOX10 strategy contained a significantly higher and heterogenous pool of oligolineage cells, such as OPCs and mature OLs (**Figure S3C and S3D**), without affecting the differentiation of the neuronal and astrocyte lineage cells, thus demonstrating that SOX10 induction in hiPSC-derived NPCs is *per se* sufficient to rapidly generate a heterogeneous pool of oligolineage cells (**Figure 2C and 2D**).

To corroborate these results, the density of MBP^+^ cells was also quantified in whole 15 μm-thick cryosections from 8-week-old organoids by immunofluorescence, showing a significant increase in in organoids stimulated with doxycycline (average of 6.4 highly branched MBP^+^ cells/organoid cryosection), compared to other experimental conditions (**Figure 2E and 2F**). Only sparse MBP^+^ cells were found in organoids not stimulated with doxycycline at week 8 (**Figure 2E and 2F**), likely due to their exposure to specific factors (*i.e.,* AA, T3) in the culture medium that can by themselves support OL differentiation and maturation. In those organoids not stimulated with doxycycline, we observed more MBP^+^ cells after 16 weeks of differentiation, suggesting that OL maturation occurs later in the absence of doxycycline stimulation (**Figure 2H, S3E, and S3F**). By electron microscopy, the quantification of myelinated axons showed either the absence or very low density of myelinated axons in doxycycline-untreated organoids compared to doxycycline-treated organoids [mean±SEM 6.4±0.9 *vs* 19.4±4.1 myelinated axons/1000 μm^2^, respectively, *p*=0.04] (**Figure 2G**).

### Organoids partially recapitulate the cell diversity of the human adult cortex

To assess the transcriptional similarity between cell types in the organoids and CNS cells of the adult *vs* fetal human brain, we mapped our single cell data onto two different reference atlases using Azimuth: (1) a single-cell RNA seq reference atlas of the adult human primary motor cortex from individuals without neurological disease (**Figure 2I**)^16,17^ and, (2) a single-cell RNA seq reference atlas of the fetal human brain (72-129 days of postconceptual□age).^18^ The mapping process allowed identification of cells with a high mapping score (>0.75) more frequently on the adult than the fetal human brain suggesting the presence of mature cells in our 3D *in vitro* platform (**Figure 2J and Figure S3G**).

Notably, the maturation stage of organoids profoundly distinguishes our organoid model from previously published cortical brain organoids.^19–26^ Cortical brain organoids are 3D cellular structures recapitulating the developing cellular architectural layering of the human cortex around a central pseudo-ventricle cavity and allowing the study of physiological and pathological processes occurring during neurodevelopment.^27^ To confirm the difference between our organoid model and cortical organoids, we re-analyzed single-cell transcriptome profiles of human cortical organoids from 8 different protocols collected from public resources^19–26^ (differentiation range ∼60/70 days; dataset metanalysis available in Tanaka et al., 2020^28^) and, similarly, we mapped them on the reference atlases of the adult *vs* fetal human brain. Of note, using a high mapping score (>0.75), in cortical brain organoids,^19–26^ only astrocytes showed the transcriptional profile of their corresponding cells in the adult cortex. We did not find any cells with a transcriptional profile of adult OLs (**Figure 2K and Figure S3H**).

In summary, these results indicate that, differently from cortical brain organoids modelling the developing brain, our multilineage organoids represent an alternative and simplified cellular platform enriched by a higher proportion of mature CNS cell types.

### Pseudotime trajectory analysis recapitulates the prominent differentiation paths for NPC-derived cells

To delineate the developmental trajectory tracing the lineage specification of NPCs in organoids as they differentiate, we used an unsupervised algorithm (Monocle) to show single-cell gene expression kinetics over time and order cells by progress through differentiation.^29^ Monocle reconstructed a trajectory capturing the progression of single cells through differentiation, which contains three termini (denoted “NEU,” “ASTRO,” and “OL”) corresponding to three different differentiation outcomes. To reach these outcomes, cells passed through two prominent branch points (denoted “1” and “2”) (**Figure 3A**), suggesting that NPCs can follow two distinct cell lineages. Cluster 7 (cycling cells) was defined as the root of the trajectory. A global differential analysis comparing the two paths away from branch point 1 indicated 115 genes with branch-dependent expression (qval=0) (**Figure 3B**) (NEU branch-associated genes *i.e., GNG3, SEZ6L2, ATP1B1, SYT1, SYT4, NSG2, SNHG14, NEFM, CALY, SNAP25, NEFL; vs* glia [ASTRO and OL] branch-associated genes *i.e., VIM, CD99, ANXA5, EDNRB, CST3, SLC1A3, AQP4, CLU, ATP1A2, GFAP, APOE, S100A16, SPARCL1*).

**Figure 3.**
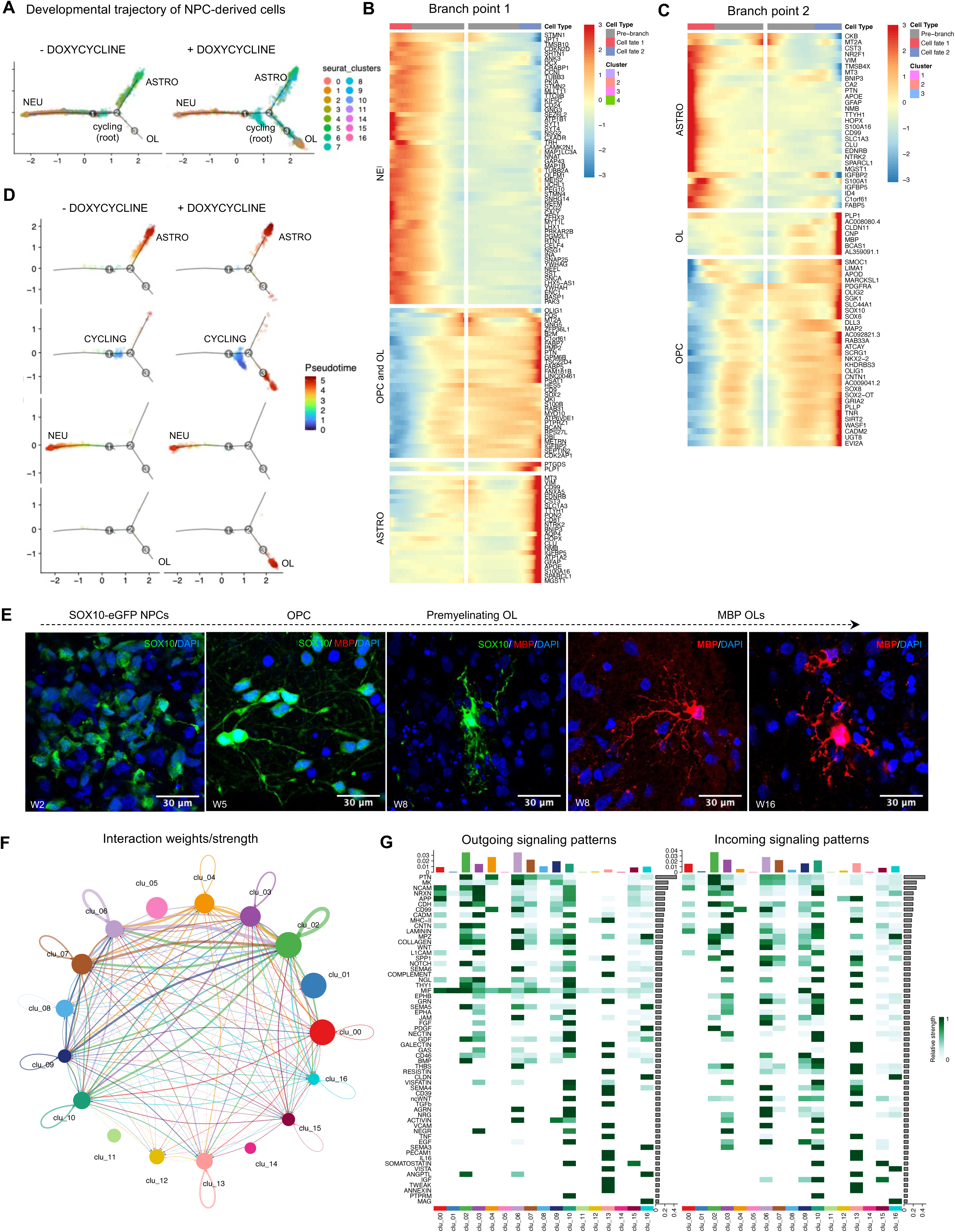
Pseudotime trajectory of the differentiation paths for NPC-derived cells and cell-to-cell communication patterns. (A) Pseudotime trajectory analysis of NPC-derived neuronal, astrocyte, and oligolineage cells, colored by Seurat cell cluster for doxycycline untreated *vs* treated organoids. (B-C) Hierarchical clustered heatmap depicting genes whose expression patterns covary across pseudotime (z-scores normalized by row) at branch points 1 and 2, respectively, for doxycycline-stimulated organoids. (D) Pseudotime trajectory analysis of selected NPC-derived neuronal (NEU) (cluster 3 and 10), astrocyte (ASTRO) (cluster 4 and 6), and OL (cluster 16), showing the high degree of their maturation, and cycling cells (CYCLING) (cluster 7), colored by pseudotime, for doxycycline untreated *vs* treated organoids. (E) Immunostaining of SOX10^+^ and MBP^+^ cells in cryosections at 2, 5, 8, and 16 weeks of differentiation in doxycycline-exposed SOX10-eGFP organoids (scale bar:30 μm). (F) Circle plot representing cellular communication among the different clusters in organoids using CellChat. Circle sizes are proportional to the number of cells in each cell group and edge width represents the communication probability. (G) Heatmap showing detailed communication through individual pathways and providing insights into the autocrine-*vs* paracrine-acting pathways for each cell type using CellChat.

Branch-dependent expression analysis at branch point 2 showed significant changes in 67 genes with ASTRO- and OL-dependent expression (ASTRO branch-associated genes *i.e., VIM, APOE, GFAP, SPARCL1, AQP4, S100A16, CD99, SLC1A3, CLU vs* OL branch-associated genes *i.e., PLP1, CLDN11, CNP, MBP, BCAS1, PDGFRA, OLIG2, SOX10, SOX6, NKX2−2, OLIG1*) (qval∼0) (qval<10^-170^) (**Figure 3C**). The developmental trajectories split by clusters show the distribution of cells within each cluster along the branches based on their differentiation stage (see representative trajectories of selected mature NEU, ASTRO, and oligolineage clusters in **Figure 3D**). Notably, the analysis revealed a spectrum of oligolineage stages ranging from OPCs expressing *PDGFRa, SOX10, SOX6, NKX2-2, OLIG2, OLIG1,* to OLs expressing myelin genes (*PLP1, MBP*) (**Figure 3C**). Consistently, we confirmed by immunofluorescence analysis the developmental progression of the hiPSC-derived oligolineage cells during the differentiation protocol: they progress from SOX10-eGFP NPCs to bipolar OPCs to premyelinating OLs to highly branched MBP^+^ OLs (**Figure 3E**). Premyelinating OLs were identified as SOX10-eGFP^+^ cells characterized by a branched morphology that did not colocalize with either MBP^+^ cells or GFAP^+^ cells, suggesting that these cells are neither myelinating OLs nor astrocytes.

By examining the highly differentially expressed genes between the two trajectories, we concluded that branch point 1 represents a decision point corresponding to whether a cell will follow a neuronal or glial differentiation route, while branch point 2 is a check point governing whether, having followed the glial lineage route, it will differentiate in either astrocyte or OLs, mimicking what is known to happen during the neurodevelopment. Of note, the ectopic *SOX10* induction by doxycycline did not alter the maturation trajectory of neurons and astrocytes, but only OLs commitment (**Figure 3A and Figure 3D**).

### Inference on cell-to-cell communication identifies patterns between neuronal, glial, and immune cells

To predict the global intercellular communication in organoids, we employed CellChat, a tool to quantitatively infer and analyze cell-cell communication networks from single-cell RNA-seq datasets, based on a signaling ligand-receptor (LR) interaction database (**Figure 3F**).^30^ CellChat analysis on single-cell RNA-seq data detected 1,138 significant ligand-receptor interactions among the 17 cell clusters, which were categorized into 142 signaling pathways. First, we explored how multiple cell populations and signaling pathways coordinate to function, by employing a pattern recognition method implemented in the “IdentifyCommunicationPatterns” function of the CellChat package. We defined 6 pattern modules for outgoing signaling and 4 signaling pattern modules for incoming communication (**Figure S4**). The choice of the number of modules is based on the suggestion provided by the silhouette plot generated by the function selectK.

Outgoing signaling pattern #2 is associated with immune cells and features a high contribution score^30^ of the MHC-II, SPP1, complement, TGFβ, TNFα, and IL-16 pathways (**Figure S4**). Out-going signaling pattern #4 includes high contribution score of signaling pathways, such as THBS, AGRN, and ACTIVIN, which characterize the astrocyte clusters. Outgoing oligolineage signaling is represented by pattern #5, including pathways such as PDGF, CLDN, SEMA3, and MAG. Outgoing neuronal signaling is defined by different pattens (#1, #6) and feature pathways such as EPHB, EPHA, FGF, NECTIN, VISFATIN, WNT, NRG, EGF, SOMATOSTATIN, and PTPRM (**Figure S4**). We performed the same analysis for incoming signaling (**Figure S4**). The pattern analysis microglial cluster signaling is dominated by pattern #2, which features a high contribution score by pathways such as APP, MHC-II, complement, THY1, CD39, and TGFβ among others. Most incoming oligolineage cells are characterized by pattern #4, which is characterized by the COLLAGEN, PDGF, CLDN, VISTA, and MAG pathways (**Figure S4**).

By cross-referencing outgoing and incoming signaling patterns, we detailed communications through individual pathways, providing insights into the autocrine-acting *vs* paracrine-acting pathways for each cell type (**Figure 3G**). In detail, major autocrine-acting pathways in immune cells are *via* the antigen presenting MHC-II complex and the complement pathway (CD46), as well as GNR, GALECTIN, the adenosine generating enzyme CD39 (regulating microglia ramification),^31^ TGFβ1^32^ (critical for adult microglia homeostasis), VCAM, PECAM1, TNFα, and IL16 signaling. Among the major paracrine-acting microglia-to-astrocyte pathways, network centrality analysis identified that microglia are the most prominent source of osteopontin (SPP1) acting not only in an autocrine fashion, but also on astrocytes (**Figure 3G**). These findings are consistent with data indicating that SPP1 exhibits cytokine, chemokine, and signal transduction functions by interacting with integrins and CD44-variant receptors, such as those expressed in astrocytes. Moreover, astrocytes and microglia were inferred to communicate also *via* thrombospondins and CD46, a regulator of complement activation, and semaphorin 4D signaling, which controls microglia and astrocyte activation (**Figure 3G**).

### Organoids respond to the exposure of MS-inflamed CSF mimicking the CNS glial-microglia phenotypes in chronic active multiple sclerosis

We tested whether our cellular model was able to respond to complex inflammatory stimuli, *i.e.*, the supernatant of inflamed cerebrospinal fluid (CSF) from subjects diagnosed with MS (**Figure 4**), and whether it could, in principle, be exploited to interrogate and decode *in vitro* the human glial-microglia axis in the context of a dysregulated inflammatory environment (simulated by the inflamed CSF).

**Figure 4.**
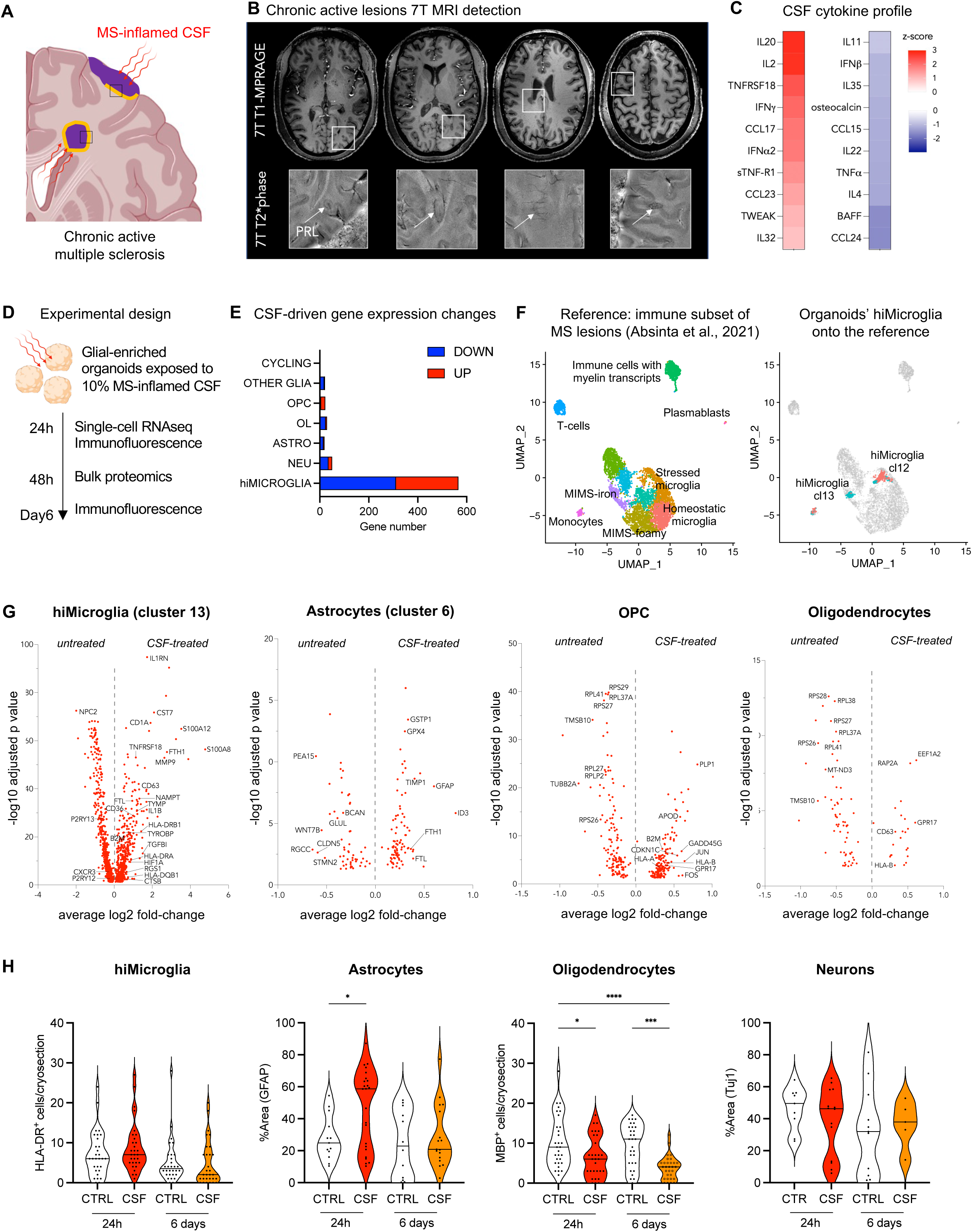
In response to inflamed CSF, organoids mimic macroglia-microglia phenotypes in chronic active multiple sclerosis. (A) Schematic of MS brain pathology with chronic active lesions (with their inflammatory edge in orange), cortical lesions and MS-inflamed CSF. (B-C) Representative case with both *in vivo* 7-tesla MRI and CSF-inflammatory profile: 54-year-old woman with relapsing MS, untreated at the time of lumbar puncture. The MRI shows 4 chronic active MS lesions seen by their paramagnetic rims (arrows, magnified views) on susceptibility-based images (B). Heatmap showing the z-scores of CSF-cytokines concentration relative to the CSF of the other 74 untreated MS cases (C). (D) Schematic of the experimental design: exposure of organoids to 10% patient-derived CSF supernatant (devoid of lymphocytes and monocytes) and processing for scRNAseq, immunostaining and bulk proteomics at different time points. (E) Bar plot of counts of significant differentially expressed genes (average log_2_ fold-change>0.5; *p* adjusted<0.05, upregulated in red and downregulated in blue) between 24-hour CSF-exposed *vs* untreated organoids by cell cluster. (F) Reference atlas based on the re-analysis of immune subclustering of chronic active MS lesions in Absinta et al.^3^ (on the left) and mapping of organoids scRNAseq data onto the MS immune subset reference (on the right). (G) Volcano plots report gene expression changes in untreated *vs* 24-hour CSF-stimulated SOX10-eGFP organoids for each glial cell population (*p* adjusted<0.05). (H) Violin plots showing the quantification of main cell populations in CSF-treated (“CSF”) *vs* untreated (“CTRL”) organoids at different time points [ANOVA test (Tukey’s multiple comparison test \**p*<0.05, \*\**p*<0.01, \*\*\**p*<0.001, \*\*\*\**p*<0.0001)]. *Abbreviations:* MS:multiple sclerosis; CSFcerebrospinal fluid; NEU:neurons; ASTRO:astrocytes; OPC:oligodendrocyte precursor cells; OL:oligodendrocytes; hiMICROGLIA:hiPSC-derived microglia; OTHER GLIA:immature astrocytes; MIMS:microglia inflamed in MS; AIMS:astrocytes inflamed in MS.

To this end, the levels of 69 inflammatory mediators were evaluated in 75 CSF samples from untreated MS cases by Bio-Plex analysis (**Table S5**). From this pool, we randomly selected CSF from 3 MS cases and additional 3 CSF from non-neurological controls (**Table S6**). We exposed organoids from 4 different iPSC lines to 10% CSF supernatant for 24 hours; RT-qPCR was then performed to measure the expression of inflammatory markers (*NAMPT, CXCL8*) and glial lineage markers (*AIF1, GFAP)* selected based on prior evidence from our snRNAseq MS study.^3^ We observed that CSF from controls did not modulate *NAMPT* and *CXCL8* mRNA expression, compared to untreated conditions, whereas treatment with MS CSF induced a significant increase in *NAMPT* and *CXCL8* mRNA expression (ANOVA *p*=0.05 and *p*=0.03, respectively; **Figure S5A**). *AIF1* and *GFAP* mRNA were not significantly different across conditions (**Figure S5A),** suggesting that the obtained results were not biased by heterogenous glial cell composition. The MS CSF-triggered inflammatory response has been also confirmed through bulk proteomic analysis conducted 48 hours after CSF exposure, revealing differential expression of 105 proteins between conditions. Pathways analysis showed enrichment in the JAK-STAT-mediated cytokines and chemokines signaling pathway (STAT3, AKT1, AKT2, AKT3, GNB2, HRAS, GNB1, GNB2, GNB4, CSK, CRKL), proteasome activation, and apoptotic processes (**Figure S5B, Table S7**).

To more deeply interrogate the effect of MS-inflamed CSF at molecular and cellular levels, organoids (*n*=30 for each experiment, 3 hiPSC lines) were exposed for 24 hours to 10% CSF supernatant and dissociated for single-cell RNA-seq (**Figure 4D**). For this experiment, we selected the CSF of an untreated 54-year-old woman with relapsing-remitting MS with chronic active lesions on her 7T MRI scan (**Figure 4B**). Although this CSF was largely representative of the inflamed

CSF of relapsing MS, we noticed z-scores>1.5 for proinflammatory cytokines and chemokines, such as IL-20, IL-2, TNFRSF8, IFNγ, TARC/CCL17, IFNα2, sTNFR1, CCL23, and TWEAK, when compared to the CSF of the other 74 untreated cases, indicating Th1-driven response (**Figure 4C**; **Table S5**). We performed the cluster-wise differential gene expression analysis between CSF-exposed *vs* nonexposed organoids. Differentially expressed genes (average log_2_ fold-change>0.5 and adjusted *p*<0.01) were implemented for enrichment analysis (enrichR package). As expected, CSF perturbation strongly affected the transcriptional profile of hiMicroglia (first responder), but also oligolineage cells and astrocytes (**Figure 4E**). On the other hand, the short exposure to CSF seems not affect neurons, with only a subset (cluster 0) shown to be enriched for ferroptosis by the pathway analysis (*MAP1LC3A*) (**Table S8**).^33^

In detail, in CSF-exposed hiMicroglia, the pathway analysis enriched for regulation of inflammatory response (*CCL22, CXCL8, CCL20, HIF1A, MMP14, SELENOS, RGS1, GPR183, IL1B, NAMPT, CD14, TIMP1, IL7R*), antigen processing and presentation (*HLA-DMA, HLA-DRA, HLA-DQA1, HLA-DRB1*), cytokine-cytokine receptor interaction (*IL1RN, CCL22, CXCL8, CCL20, IL1B, TNFRSF18, CCL4, CCL3, IL7R, IL2RG, TNFRSF4*), activation of toll-like receptor signaling pathway (*CXCL8, IL1B, CCL4, CCL3, MAP3K8, CD14*) complement pathway (*FCN1, FCER1G, HSPA5, MMP14, S100A12, ADAM9, TIMP1, CD36, CALM1, S100A9, CTSB*), reactive oxygen species production (SELENOS, NDUFB4, PRDX1, CAT, GLRX), ferroptosis (*FTH1, ACSL4, FTL*), and interferon g response (*NAMPT, FPR1, ITGB7, PTPN6, IFI30, HIF1A, HLA-DQA1, HLA-DRB1*) (**Figure 4G**, **Table S8**).

In oligolineage cells (both OLs and OPCs), we found a significant downregulation of ribosomal genes, suggesting modifications of the cellular translational machinery. Of note, ribosome stalling during protein synthesis may harbor proteotoxic components and trigger cellular stress conditions, as observed during neurodegeneration.^34^ Moreover, upregulation of *GPR17,* a known negative regulator of myelination recently investigated in MS,^35^ was observed (**Figure 4G**). Upregulation of the tetraspanin CD63 might indicate release of extracellular vesicles by the OLs in response to inflammatory insult. In OPC (cluster 2), treatment with CSF showed enrichment for genes implicated in TNFα signaling *via* NF-kB (*JUN, FOS, IER2*), as well as in hypoxia (*CDKN1C, JUN, FOS*), IL-2/STAT5 (*CDKN1C*, *ALCAM*), and p53 (*JUN*, *FOS*) signaling pathways (**Figure 4G**, **Table S8)**.

Next, to assess transcriptional similarity between the cell types in the CSF-exposed organoids and cells in the MS brain, we mapped our single cell data onto the single-nucleus RNAseq dataset from autopsy MS brain tissue in Absinta et al., 2021^3^, which included chronic active MS lesions (**Figure 4F**). Notably, the transcriptional profile of hiMicroglia partially overlapped with the signature of microglia inflamed in MS (MIMS)-iron of the chronic active lesion edge^3^ (**Figure 4F**). Of interest, overlapping genes include the upregulation of iron-related genes (*FTL, FTH1*) — consistent with the iron retention within microglia observed by MRI at the chronic active lesion edge (PRL imaging biomarker, **Figure 4B**) — as well as immunoglobulin Fc-gamma receptors (*FCGRT*), MHC class II protein complex (*HLA-DRA*), *SPP1,* and the inflammatory cytokine *IL1B*.^3^ We also observed downregulation of homeostatic microglial genes, such as *CX3CR1* and *P2RY12*.

To quantify such similarity between MIMS-iron and CSF-perturbed hiMicroglia, we scored their transcriptomic profile by gene set enrichment analysis (GSEA). Interestingly, we identified a significant enrichment score for both upregulated (cluster 12: normalized enrichment score [NES]=1.69, adjusted *p*<0.01; leading edges: *FTH1, CTSB, CD63, FCER1G, B2M, SPP1, ITGB2, APOE, HIF1A, GRN, TYROBP, CTSZ, TIMP2, CD9;* cluster 13: NES=1.60, adjusted *p*< 0.04, leading edges: *FTH1, FCER1G, ITGB2, CD63, HIF1A, CSTB, SPP1, TYROBP, CTSS, CTSZ*) and downregulated signatures (cluster 13: NES=-1.67, adjusted *p*<0.03; leading edges: *CX3CR1, MEF2A, FCHSD2, P2RY12, CYTH4*) (**Figure S5C**), providing additional statistical evidence of transcriptional profile similarity.

A similar approach was implemented for the astrocytes, where we found that the transcriptional profile of inflamed astrocytes partially overlapped with the signature of designated “astrocytes inflamed in MS” (AIMS)^3^ and reactive astrocytes at the chronic active lesion edge (**Figure S5D and S5E**). This indicates that the astrocytes in organoids can acquire disease-associated phenotypes providing a simplified model for their investigation.^3^ Interestingly, in the comparison between controls-vs MS-derived cells, the CSF challenge was differentially enriching the MS-astrocytes and MS-neurons in energy-related terms (*i.e.,* mitochondrial electron transport, oxidative phosphorylation, cellular response to hypoxia) [**Table S9**].

Similarly, organoids’ oligolineage cells co-map with subclusters of OPC and early myelinating OL seen in the MS tissue (**Figure S5F and S5G**).^3^ As highlighted in recent human snRNAseq MS studies,^3,36^ we found some OL heterogeneity in our organoid model: 17.8% of them mapped on cluster 2 of Absinta et al.^3^ (*OPALIN*^+^ as described also in Jäkel et al.^36^) and the remaining 82.2% on cluster 6 (*BCAS1+* suggesting they are newly myelinating oligodendrocytes) of Absinta et al.^3^

### Cell-to-cell communication perturbation by MS-inflamed CSF

Using CellChat, we observed differences in the number of interactions (defined as the delta of the number of ligand-receptor pairs per connection) and interaction strength (defined as the delta of the sum of interaction weights of ligand-receptor pairs per connection) between CSF-treated and untreated organoids (**Figure S6A**). In the CSF-treated organoids, there was a higher number of interactions between iMicroglia and astrocytes, while the interaction strength was greater between astrocytes and neurons (shown as red lines in **Figure S6A**). Some crosstalk, like the OPC-astrocytes interaction, displayed fewer interactions but higher interaction strength, suggesting a complex remodeling of communication patterns. Upon detailed exploration of the upregulated lig-and-receptor pairs in CSF-treated organoids (**Figure S6B**), we noted an upregulation in microglial MHCII-signaling and an increase in microglial-osteopontin (*SPP1*) interaction. *SPP1* interaction was not limited to integrins (*ITGAV* and *ITGB1*) with both astrocytes and oligolineage cells but also involved *CD44* through an autocrine pattern (upregulation of both ligand and receptor in microglial cells exposed to the CSF). These findings support and expand upon our results from the differential expression analysis and aligns with observations from our snRNAseq analysis of chronic active MS brains.^3^ It is worth noting that exposure to inflamed CSF increased the overall signaling of neuronal and glial *NCAM1*, which plays a crucial role in CNS processes like cell adhesion, migration, synaptic plasticity, and the regulation of inflammatory signaling through NFkB and MAPK pathways.^37^

### Short and long-term effects of MS-inflamed CSF on the glial-enriched organoids

The effect of MS-inflamed CSF on the organoid cultures was examined after 24 hours and 6 days using different experimental approaches to fully understand its temporal dynamics. Beside the early transcriptomic changes across various cell populations, we further investigated whether CSF exposure could lead to delayed oligodendrocyte and neuronal loss. By immunostaining, we observed a significant decrease (nearly 50%) in the number of MBP^+^ OLs (ANOVA p<0.0001), but not in neuronal staining area 6 days following the inflamed MS-CSF exposure, suggesting specific vulnerability of OLs to the MS-CSF (**Figure 4H**). The count of HLADR^+^ (antigen-presenting) microglial cells and astrogliosis (GFAP staining area) did not exhibit a significant difference between control and CSF-treated organoids at day 6, but only a transient increase of GFAP staining after 24 hours (p=0.004; **Figure 4H**).

## DISCUSSION

In the present study, the obtained results serve two purposes. The first part of the study introduced a protocol for generating a 3D hiPSC-based organoid model, partially able to recapitulate the rich diversity of cell types in the *in vivo* human brain. This model offers some relevant advantages over existing methods, including the relatively rapid generation of mature CNS cell types, an enrichment of glial cells (including MBP^+^ OLs), and the integration of hiPSC-derived microglia into organoids. By short ectopic *SOX10* expression, we were able to promote oligodendroglia commitment in a subset of NPCs, without disrupting the oligodendroglia endogenous differentiation program as well as the differentiation of other NPC-derived cells, such as neurons and astrocytes. In the second part of the study, the organoid model was applied in a specific translational context. Due to the absence of preclinical models for smoldering MS, we exposed the glial-enriched organoids to inflamed CSF supernatant from MS patients. This approach aimed to model and to explore the spatial and temporal interactions between macroglia-microglia, which are considered crucial to the non-relapsing biology of progressive MS.

Differently from cortical brain organoids modelling the developing brain, our model might represent a simplified platform lacking cortical patterning but enriched by mature glial cell types. In previously published cortical brain organoid single-cell transcriptomic datasets,^19–26^ only astrocytes showed the transcriptional profile of their corresponding cells in the adult cortex, and no cells with a transcriptional profile of adult OLs were detected. Of relevance, immunohistochemical analysis of the human fetal brain demonstrated early OL specification^38–42^ and myelination beginning prenatally.^43^ The overall lack of OLs in cortical organoids likely prevents a proper modeling of the developing brain in terms of the spatial and temporal progression of oligolineage, as well as of the regulatory signals for OL differentiation.^44^ To date, only three other studies generated hiPSC-derived brain organoids containing OLs (termed human OL spheroids) by adding specific small molecules (platelet-derived growth factor-AA, insulin-like growth factor-1, 3,3’,5-triiodothronine) known to foster oligodendrocytes induction and maturation.^22,45,46^ However, compared to the Madhavan et al. and the Marton et al. protocols^22,45^ requiring 14-20 weeks of differentiation to obtain MBP^+^ OLs, our protocol offers faster and more robust production of OLs (8 weeks of differentiation) by genetically inducing the commitment of a subset of NPCs to oligodendrogenesis. Moreover, the immune counterpart is missing in the abovementioned protocols. Interestingly, we observed the presence of BCAS1^+^ OLs marking newly myelinating OLs, which have been described not only in the fetal and early postnatal human white matter, but also in a proportion of chronic MS lesions with signs of remyelination.^47^ Thus, mapping ongoing myelin formation, the presence of BCAS1^+^ OLs offers the possibility to investigate potential cellular targets for remyelination therapies in MS.

In MS, growing evidence are suggesting that the biology behind disease progression independent of relapse activity (often termed PIRA or silent progression) involves compartmentalized CNS inflammation and neurodegeneration rather than peripheral lymphocytes attacks on the CNS.^2^ In this context, it has been hypothesized that meningeal inflammation initiates a “surface-in” gradient of tissue injury implicating soluble CSF inflammatory molecules in triggering microglia and astrocytes priming as well as neurodegeneration.^48,49,50^ To experimentally test this hypothesis and to recapitulate such pathological processes, including their temporal sequence, in this study, organoids were exposed to the inflamed CSF of MS patients with chronic active lesions at different time points (from 24 hours to one week).

Short-term MS-inflamed CSF perturbation strongly affected the transcriptional profile of hiMicroglia (acting as first responder), oligolineage cells, and astrocytes, but not neurons, indicating that CSF-induced inflammation may drive neurodegeneration by first triggering glial changes.^33^ The transcriptional profile of hiMicroglia partially overlapped with the signature of MIMS-iron at the chronic active lesion edge.^3^ In line with our experimental findings, a recent human pathological study highlighted the correlation between meningeal inflammation and higher proportion of chronic active lesions in the MS brain.^51^ The possibility to mimic the activation of microglia genes modules implicated in antigen presentation, direct propagation of inflammatory responses, and oxidative damage in the context of a 3D human cellular environment is encouraging. However, no upregulation of phagocytic activity, as seen in the MIMS-foamy of the chronic active MS lesions,^3^ was observed in the CSF-exposed hiMicroglia, potentially supporting divergent pathways of activation. We also demonstrated that CSF-exposed astrocytes in organoids can acquire the astrocyte phenotype of both reactive and inflamed astrocytes (AIMS) seen at the chronic active lesion edge, thereby providing a simplified model for their investigation. In OPCs, CSF induced the upregulation of genes involved in self-antigen presentation, a process recently observed after OPC exposure to IFNγ in both cell cultures and animal MS models.^52,53^ Together with downregulation of the cellular translational machinery, such evidence support the vulnerability of both OLs and OPCs in the MS inflammatory environment, as further demonstrated by nearly 50% reduction in OLs, but not neurons by day 6. Our temporally resolved organoid data support and expand on the role of soluble CSF mediators in sustaining a downstream cascade of events leading to oligodendrocytes death and potentially neurodegeneration.

Notably, there was a distinct response observed in organoids derived from individuals with MS compared to those from the control group when exposed to the MS-inflamed CSF. By single-cell transcriptomics, an enrichment in terms associated with cellular energy (such as mitochondrial electron transport, oxidative phosphorylation, and cellular response to hypoxia) was observed in MS-derived organoids, particularly within MS-neurons and MS-astrocytes (**Table S9**). These findings align with the concept of higher energy demand of MS brains as well as with prior studies investigating intrinsic differences between hiPSC-derived cells from MS patients and control subjects.^14,15,54^ Furthermore, supporting this observation, the recent effort of the International Multiple Sclerosis Genetics Consortium underscored that genetic variants linked to disease susceptibility map on immune cells, whereas disease severity seems to be linked to CNS cells.^55^ This suggests some potential genetic determination of the dysfunctional CNS response to inflammatory injury in MS.

In conclusion, our *in vitro* cellular platform provides an important early foundation with which to iteratively increase our understanding by testing hypotheses emerging from human association studies. The scalability of hiPSC-derived organoids also makes this system amenable to genetic and small molecule screens to identify novel therapeutics and unveil the key regulators of human myelination and chronic inflammatory neurodegeneration.

## LIMITATIONS OF THE STUDY

Our 3D model represents a simplified platform disentangling the complexity of the cellular landscape of the human brain, and it does not entirely recapitulate the vast amount of its cellular heterogeneity (*i.e.,* vascular cells are completely lacking) as well as its complex cytoarchitecture. Moreover, although we provided a thorough characterization of our model, the experiments were limited to 5 different cell lines derived from 2 non-neurological controls and 3 MS cases; thus, the number of donors could be increased, especially to assess morphological and transcriptional differences between MS and controls. To overcome inter-organoid variability limitations, next steps might also include the generation of multi-donor organoids. Finally, since microglia can survive in culture for only few weeks once mature, further optimization of the protocols would be necessary to establish a more chronically inflamed organoid model.

## ACKNOWLEDGMENTS

This work has been supported by Cariplo Foundation (2019-1677), Fondazione Regionale per la Ricerca Biomedica (FRRB) Early Career Award (1750327), Roche Foundation for Independent Research 2019, Adelson Medical Research Foundation, International Progressive MS Alliance (PA-1604-08492 BRAVEinMS), and Intramural Program of the NINDS, NIH.

We thank the members of the Neuroimmunology Clinic (led by Irene Cortese), Viral Immunology Section (led by Steven Jacobson), and Translational Neuroradiology Section (led by Daniel Reich) of NINDS,NIH for CSF collection and 7T MRI acquisition; Luigi Naldini’s group (TIGET) for providing plasmids; Maira Gironi for recruiting patients for skin biopsy; Annamaria Cafarella for generating the lentiviral vector; Alessandra Mandelli, Maria Sofia Martire, Svetlana Bezukladova, Carolina Peri, and Jessica Perego for helping with cell cultures and/or FACS; Giacomo Sferruzza, Giovanni Landoni and Antonella Crescenti for CSF collection from non-neurological controls; Francesca Giannese and Caterina Oneto (COSR) for helping with the 10X Genomics experiments; Valeria Berno and the resources of the ALEMBIC facility for confocal imaging; resources of the Research Cluster of San Raffaele Scientific Institute for computational analysis.

## AUTHOR CONTRIBUTIONS

Conceptualization: MA; methodology: FF, EP, ST, EB, FR, TK, VM, PP, AQ, AAP, CB, LS, DSR, PPB; writing original draft: FF, MA; intellectual contribution and manuscript editing: MA, LS, TK, PAC, DSR, PPB, GM; funding acquisition and supervision: MA.

## STAR METHODS

### RESOURCE AVAILABILITY

#### Lead contact

Further information and requests for resources and reagents should be directed to and will be fulfilled by the lead contact, Martina Absinta.

#### Materials availability

All stable reagents generated in this study are available from the lead contact without restriction except for human induced pluripotent stem cell lines and their derivative for which permission must be requested and a material transfer agreement must be completed.

#### Data and code availability

- Single-cell RNA-seq data have been deposited at GEO (GSE233295) and are publicly available as of the date of publication. Microscopy data reported in this paper can be shared by the lead contact upon request.
- All original code has been posted in GitHub (https://github.com/AbsintaLab/GLIA-ENRICHED-ORGANOIDS).
- This paper analyzes existing, publicly available genomic datasets. These accession numbers for the datasets are listed in the key resources table.
- Any additional information required to reanalyze the data reported in this paper is available from the lead contact upon request.

### EXPERIMENTAL MODEL AND SUBJECT DETAILS

#### Ethics statement

The study was approved by the ethical committee of San Raffaele Hospital, Milan, Italy (45/INT/2021 and 137/2021).

#### Human subjects

Permission to generate human induced-pluripotent stem cells (hiPSC) from human fibroblasts was granted by the ethical committee of the San Raffaele Hospital (Milan, Italy) (BancaINSpe - INSPE1178). Fibroblasts were isolated from skin biopsies from 2 healthy volunteers (2 women) and 3 individuals with a diagnosis of relapsing-remitting MS (2 women, 1 man) (see **Table S1** for demographics). MRI acquisition and collection of CSF by lumbar puncture from 4 MS cases (2 women, 2 men) (see **Table S6** for demographics) were approved by the National Institute of Health (Bethesda, US) (protocol number: 89-N-0045; principal investigator: Daniel S. Reich, MD, PhD) and shared under a materials transfer agreement. CSF collection by lumbar puncture from 3 non-neurological controls (1 woman, 2 men) was granted by the ethical committee of the San Raffaele Hospital (Milan, Italy) (BancaINSpe - INSPE1178) (see **Table S6** for demographics). Written consent was obtained from all subjects.

#### Human induced-pluripotent stem cells derived NPCs

Fibroblasts were reprogrammed into iPSCs by using the replication-incompetent Sendai virus kit Cytotune (Invitrogen), according to manufacturer’s instructions. To generate NPCs according to Reinhardt *et al,*^56^ colonies of iPSCs were detached from mouse embryonic fibroblasts by treatment with 1 mg/mL collagenase IV. After sedimentation, cells were resuspended in human embryonic stem cell (hESC) medium (DMEM-12, 20% KO serum replacement, 1:100 non-essential AA, 1:100 penicillin/streptomycin, 1:100 L-glutamine, 1:200 di β-mercaptoethanol), supplemented with 1 μM dorsomorphin (Tocris), 3 μM CHIR99021 (Tocris), 10 μM SB-431542 (Ascent Scientific) and 0.5 μM purmorphamine (Enzo). Embryoid bodies (EBs) were formed by culturing cells in non-tissue culture, extra low adhesion petri dishes. On day 2 medium was changed to N2B27 medium containing equal parts of neurobasal (GIBCO) and DMEM-F12 medium (Gibco) with 1:100 B27 supplement lacking vitamin A (GIBCO), 1:200 N2 supplement (GIBCO), 1% penicillin/streptomycin/glutamine (PSG) and the same small molecules mentioned before. On day 4, dorsomorphin and SB-431542 were withdrawn, while ascorbic acid (AA; 150 μM) was added to the medium. On day 6, EBs were cut into smaller pieces and plated onto matrigel (Matrigel Growth-factor-reduced, Corning) coated 12-well plates. For passaging, NPCs were treated with Accumax. After 3 passages purmorphamine was replaced by 0.5 μM SAG (NPC medium).

## METHODS DETAILS

### Lentiviral Vector Construction

To stimulate the production of OLs in hiPSC-derived NPCs, we generated a lentiviral vector encoding the human SOX10 coding sequence and green fluorescent protein (GFP) as reporter gene under the control of doxycycline-inducible promoter (**Figure S1A**).

### Enzymatic modification and DNA ligation

To obtain suitable fragments for cloning, enzymatic restriction was performed digesting the plasmids with the appropriate restriction enzymes. #945.pCCL.sin.cPPT.SV40ployA.eGFP.minCMV.hPGK.deltaLNGFR.Wpre plasmid (kindly provided by Prof. Luigi Naldini group, San Raffaele Telethon Institute for Gene Therapy, Milan) was digested with the XbaI and NheI restriction enzymes to obtain the vector, and the plasmid #115241 - pZ M2rtTA_CAGG_TetON-Sox10 _GFP (Addgene) was digested with MluI and SpeI restriction enzymes to obtain the insert. When needed, the DNA was filled in with Klenow (New England Biolabs) or T4 DNA Polimerase (New England Biolabs) to obtain blunt ends. The linearized vector plasmid was dephosphorylated by Antarctic Phosphatase (AnP) (New England Biolabs. The ligation between the insert and the linearized plasmid (vector) was performed with T4 DNA-ligase (New England Biolabs). Then, the ligation product was used to transform chemically competent bacteria.

### Bacterial transformation

Chemically competent bacteria (One Shot TOP10; Cat. No. C404010, Invitrogen) were used. Bacteria were stored at -80°C, ready to use. The bacteria were thawed in ice and 10 μL of ligation was added to each vial and incubated for 30 minutes on ice and, subsequently, heat-shocked at 42° C for 30 seconds and chilled on ice for 2 min. Then, 250 μL of pre-warmed S.O.C. medium (Invitrogen) were added to the bacteria plus plasmidic DNA mix and shaken at 37°C for 1 hour with vigorous shaking. At the end of the incubation, the bacteria were plated in a LB-Agar plate with an antibiotic resistance, *i.e.,* 100 mg/mL ampicillin (Ampicillin, Roche). The plates were incubated at 37°C overnight. For colony selection and expansion of the cultures, several colonies were picked and inoculated in LB medium with 100 μg/mL ampicillin at 37°C with shaking overnight. Then, the plasmid DNA from cultures was isolated using a Plasmid DNA Extraction mini kit (Fisher Molecular Biology, Cat. No. DE-034), according to the manufacturer’s instructions.

To obtain a high quantity of DNA, a maxi-extraction of DNA was performed using a commercially available kit (Xtra Maxi EF, Macherey-Nagel; Cat. No. 740424.50), according to the manufacturer protocols.

### Viral production and virus titration

Human Embryonic Kidney (HEK293T) cells were used for both the viral production and virus titration. HEK293T were cultures at 37°C and in 5% CO_2_. The HEK293T cells were grown in Iscove’s modified Dulbecco medium (IMDM, Sigma) supplemented with 10% of Fetal Bovine Serum (FBS, Cambrex), Ultra-glutamine (2 mM, GIBCO) and a combination of antibiotics (100 U/mL penicillin/streptomycin, GIBCO).

### Transfection and viral production

We used a third-generation *sin* lentiviral vector, produced by transient transfection into HEK293T cells, as previously described by Follenzi et al.^57^ Approximately 24 hours before the transfection 9 × 10^6^ HEK293T cells/dish were plated in a final volume of 20 mL of complete IMDM and the medium was changed with fresh new medium 2 hours before the transfection. The cells were transfected with the three packaging plasmids, the plasmid coding for the envelope protein VSVG, the plasmid coding for the packaging proteins MDLg/pRRE (which contains only the gag and pol coding sequences from HIV-1 and 374-bp RRE containing sequence from HIV-1 immediately down-stream of the pol coding sequences), the plasmid coding for REV protein and the transfer vector #945_hSox10 using CaCl_2_ and HBS pH 7.14 solution. The conditioned medium was collected after further 30 hours and filtered through 0.22 μm pore-size cellulose acetate filters (Millipore). The supernatant was centrifuged for 2 hours at 20000 rpm at 20°C. The pellet was suspended with 70 μL of 1X Phosphate Saline Buffer (PBS). The pellet suspended was left for 30 minutes at room temperature (RT). Then, the virus was harvested and shaken for 30 minutes at 4°C to obtain a homogenous solution. The viral solution was aliquoted and stored at –80°C.

### Infection of HEK293T cells and virus titration

Before viral titration, each virus was frozen and thawed once. Serial dilutions ranging from 1:10^3^ to 1:10^7^ of the virus were used to infect HEK293T cells. After 15 days *in vitro*, the HEK293T cells were harvested, then the DNA was extracted and analyzed by a real-time quantitative PCR (QuantStudio3, ThermoFischer), using the following primers: *GAPDH* #Hs00483111_cn as housekeeping gene; for the sequence of LV we used Probe LV (5’-ATCTCTCTCCTTCTAGCCTC-3’) and Primer FW (5’-TACTGACGCTCTCGCACC-3’), Primer REV (5’-TCTCGACGCAGGACTCG-3’) (ThermoFisher). The viral titer was 2 × 10^7^ Transducing Units (T.U.)/mL.

### NPC infection and sorting of GFP^+^ cells

Then, hiPSC-derived NPCs were seeded at a confluence of 10 × 10^6^ cells in T25 flask (day –1). At day 0 and 4, NPCs were infected with the SOX10-eGFP-expressing lentivirus (MOI=0,06). At day 7, NPCs were stimulated with 1 μg/mL doxycycline for 24 hours, and, then, infected cells with a stable infection constructs were selected by FACS-gating of GFP^+^ cells, using a high-speed flow cytometry sorter (Beckman Coulter MoFlo Astrios EQ).

### Production of multi-lineage organoids

To generate self-organizing forebrain SOX10-eGFP organoids^4^, we leveraged a well-characterized approach.^6,58^ NPCs were expanded in Matrigel-coated flasks in N2B27 medium containing equal parts of Neurobasal (Invitrogen) and DMEM-F12 medium (Invitrogen) with 1:100 B27 supplement lacking vitamin A (Invitrogen), 1:200 N2 supplement (Invitrogen), 1% penicillin/streptomycin/glutamine, supplemented with 3 μm CHIR, 1 μM SAG, 150 μM ascorbic acid. At 100% confluence, NPCs were enzymatically detached and counted. 2,5 × 10^6^ cells per well were plated in 3 mL of Reinhardt complete medium in non-treated 6 well-plates. Cells were grown in Reinhardt complete medium for 24 hours under constant gyratory shaking (88 rpm). Subsequently, medium was changed to glial induction medium (GIM) (100% DMEM-F12, 1:100 B27 lacking vitamin A, 1:200 N2, 1% Penicillin/Streptomycin, and 1% L-Glutaminase), supplemented with 10 ng/mL triiodo-L-thyronine (T3), 10 ng/mL rhPDGF, 10 ng/mL rhNT3, 1 μM SAG, 10 ng/mL μM IGF-1, 200 μM ascorbic acid, and 1 μg/mL doxycycline from day 1 to day 3. From day 5 till the end of the differentiation protocol, medium was replaced with glial differentiation medium (GDM) (100% DMEM-F12, 1:100 B27 lacking vitamin A, 1:200 N2, 1% Penicillin/Streptomycin, and 1% L-Glutaminase), supplemented with 60 ng/mL T3, 10 ng/mL rhNT3, 10 ng/mL IGF-1, 200 μM ascorbic acid, and 100 μM cyclic AMP analog (dbcAMP) to promote OPC proliferation and maturation. Doxycycline at the final concentration of 1 μg/mL was added to GDM medium from day 5 until day 4. On day 14, 0.01 μg/mL human recombinant GDNF and BDNF were added to the culture medium till the end of the differentiation protocol. Half of the medium was changed three times a week. Cultures were maintained at 37°C, 5% CO_2_ under constant gyratory shaking (88 rpm) for up to 8 weeks.

### Production of organoids using GIBCO protocol

To generate self-aggregating GIBCO organoids, 2 × 10^5^ hiPSCs were seeded as small colonies per well in a matrigel-coated 6-well plate in mTeSR^TM^ Plus medium, supplemented with 10 μM Y-27632. The next day, medium was replaced with neural induction medium (Neurobasal medium and 1X neural induction supplement). After 7 days, NPC colonies were dissociated and expanded in the Neural Expansion medium (50% Neurobasal, 50% Advanced DMEM/F12 media, 1X Neural induction supplement, and 5 μM Y-27632). Cultures were kept at 37°C, with 5% CO_2_ and 5% O_2_, with every second-day medium change. Organoids were generated as for Pamies et al., 2017.^6^

Briefly, 2 × 10^6^ NPCs per well were plated in uncoated 6-well plates. After 2 days, Neural Expansion medium was replaced with differentiation medium (B-27^TM^ Electrophysiology kit, 1% L-Glutaminase, 0.01 μg/mL human recombinant GDNF, 0.01 μg/mL human recombinant BDNF, 1% Pen/Strep). Cultures were kept at 37°C, 5% CO_2_, and 20% O_2_ under constant gyratory shaking (88 rpm, 19 mm orbit) for 8 weeks. Half of the medium was changed three times a week.

### hiMicroglia differentiation

Parallelly, we differentiated hiPSC from donor 1 (**Table S1**) into microglia. hiPSC were first differentiated into hematopoietic progenitors’ cells using STEMdiff^TM^ Hematopoietic Kit (Catalog #05310), and, then, STEMdiff^TM^ Microglia Differentiation Kit (Catalog #100-0019), and STEMdiff^TM^ Microglia Maturation Kit (Catalog #100-0020) were used to generate mature microglia, according to the manufacturer’s instructions.^12^ FACS analysis was performed to check each maturation stage, according to the manufacturer’s instructions.^12^ At day 38 of differentiation, microglia was added to 8-week-old organoids (to have 10% microglia into the neuron-glia network, 200,000 cells/well of a 6-well-plate were plated). Organoids were harvested 24 or 48 hours later for down-stream analysis.

### Cryopreservation

Organoids were fixed in 4% paraformaldehyde (PFA) for 1 hour at room temperature and, then, washed in 1X PBS (3 times) and transferred to 30% sucrose solution overnight. Subsequently, they were embedded in OCT and stored at −80 °C. For immunofluorescence analysis, 15-μm-thick sections were cut using a Leica cryostat.

### Immunofluorescence on whole cryosections

Cryosections were air dried for at least 40 min and, then, rinsed twice with 1X PBS for 5 min. Non-specific binding sites were blocked by incubation in the blocking solution (1X PBS containing 0.1% Triton X-100 (Sigma-Aldrich, X-100), 10% serum) for 1 hour at RT. Cryosections were incubated at 4°C for 72 hours with primary antibodies diluted in blocking solution in a humidity chamber. The following primary antibodies were used for immunofluorescence: anti-GFP (rabbit, 1:500, Life Technologies, A11122); anti-Nestin (mouse, 1:500, Millipore, MAB5326); anti-PAX6 (rabbit, 1:500, Biolegend, 901301); anti-GFAP53 (rabbit, 1:500, Agilent DAKO, Z0334); anti-S100β (rabbit, 1:300, DAKO, Z0311); anti-PDGFRα (rabbit, 1:100, Cell Signaling, 5241); anti-BCAS1 (mouse, 1:100, Santa cruz, sc-136342); anti-MBP (rat, 1:50, Millipore, MAB386); anti-Human CD45 (mouse, 1:500, Dako, M0855); anti-CD11b (rat, 1:100, Biorad, MCA74GA); anti-IBA1 (rabbit, 1:500, Wako 019-19741); anti-TMEM119 (rabbit, 1:500, Sigma, HPA051870. For double labeling, some primary antibodies were incubated simultaneously. After incubation with the primary antibodies, cryosections were washed three times for 5 min with 1X PBS containing 0.1% Triton X-100. Then, cryosections were incubated for 2 hours at RT in the dark with appropriate fluorophore-conjugated secondary antibodies (Alexa Fluor 488, Alexa Fluor 546, and Alexa Fluor 633), diluted in 1X PBS containing 1% serum and 0.1% Triton X-100. In all immunofluorescence staining, nuclei were stained with DAPI (1:25,000; Roche) in 1X PBS for 2 min. Sections were mounted with Dako fluorescent mounting medium and imaged on Olympus FluoVIEW FV3000RS confocal microscope, or Maving RS-G4 Confocal. Images were processed in ImageJ (Fiji Version 2.0.0).

### Immunofluorescence on free floating organoids

Organoids were fixed in 4% PFA for 1 hour at RT, and washed with 1X ice-cold PBS (3 times/10 minutes each) at RT. Then, fixed organoids were washed by rinsing with DPBS-Tween (0.1% Tween-20 in 1X DPBS) for 10 min at 60 rpm on the elliptical shaker at RT (3 times). For permeabilization and blocking, organoids were resuspended in blocking buffer (1% BSA and 2% Triton X-100 in 1X DPBS) and incubated for 4 hours at RT on a rocker at 80 rpm. Then, organoids were incubated with selected primary antibody on a rocker at 80 rpm for 72 hours at 4°C. The following primary antibodies were used for immunofluorescence: anti-GFP (rabbit, 1:500, Life Technologies, A11122); anti-Nestin (mouse, 1:200, Millipore, MAB5326); anti-Tuj1 (mouse, 1:1000, Biolegend, 801202). After primary antibody labeling, organoids were rinsed with wash buffer for 30 min on a rocker at 80 rpm (3 times). For secondary antibody labeling, organoids were incubated with secondary antibody at 4°C on a rocker at 80 rpm for 24 hours, protected from light (see above). Then, organoids were rinsed with wash buffer and placed in a rocker at 80 rpm for 10 min (3 times). For nuclei staining, organoids were incubated with DAPI (1:10000; Roche) diluted in 1X cold PBS and incubated for 2 hours on a shaker in dark and rinsed with buffer on a rocker at 80 rpm for 30 min (3 times). For organoids preparation for 3D Imaging, organoids were transferred into an Eppendorf tube and incubated in RapiClear^TM^ per well for a 96-well plate (F-bottom, clear, black, CellStar, Cat.No #655090) for 24 hours and, then, processed for confocal imaging.

### Transmission electronic microscopy

Organoids were incubated with Karnovsky’s fixative (2% paraformaldehyde and 2.5% glutaraldehyde). Then, the organoids were washed in 0.1M cacodylate buffer. Post-fixed in 2% osmium, 1,5% potassium ferrocyanide for 1 hour on ice, rinsed, dehydrated, and embedded in Epon resin (Electron Microscopy Science). Ultrathin sections were mounted onto slot grids for viewing using Talos 120C (Fei) microscope, images were acquired with a 4k x 4k Ceta CMOS camera (Thermo Fisher Scientific).

### Real-time quantitative PCR (qPCR)

qRT-PCR was performed in NPC and organoids using predeveloped TaqMan Assay Reagents on an ABI Prism 7500 Sequence Detection System (Applied Biosystems), according to manufacturer’s protocol. RNA was extracted using the RNeasy Mini kit (Qiagen, 74106) according to the manufacturer’s instructions. Then, RNA was eluted from columns using 20 μL of RNase-free water, and their concentrations were determined spectro-photometrically by A260 (Nanodrop-ND1000). cDNA was synthesized from 100 to 500 ng of total RNA using the Thermo Script RT-PCR (Invitrogen, 11146-024). Approximately 25 ng of cDNA was used for RT-qPCR using predesigned TaqMan Gene Expression Assays (Applied Biosystems) on an ABI Prism 7500 Sequence Detection System (Applied Biosystems). GAPDH was used as housekeeping gene. Primers used are listed in Resource Table (**Table S10**).

### Electrophysiology

Organoids were plated in GDM on glass slides (pre-coated with 15 μg/mL polyornithine in 1X PBS at 37°C, followed by incubation with 5 μg/mL laminin in 1X PBS for at least 2 hours at 37°C) for 2 days. Whole-cell patch clamp recordings were performed in cells from organoids cultured on glass slides (diameter: 1 cm) between week 8 and 12 of culture to verify the presence of neuronal electric activity. Individual slides were submerged in a recording chamber mounted on the stage of an upright BX51WI microscope (Olympus, Japan) and perfused with artificial cerebrospinal fluid (ACSF) containing (in mM): 125 NaCl, 3.5 KCl, 1.25 NaH_2_PO_4_, 2 CaCl_2_, 25 NaHCO_3_, 1 MgCl_2_, and 11 D-glucose, saturated with 95% O_2_ 5% CO_2_ (pH 7.3) at RT. Whole-cell current- and voltage-clamp recordings were performed using borosilicate pipettes filled with a solution containing the following (in mM): 30 KH_2_PO_4_, 100 KCl, 2 MgCl_2_, 10 NaCl, 10 HEPES, 0.5 EGTA, 2 Na_2_-ATP, 0.02 Na-GTP, (pH 7.2 adjusted with KOH, tip resistance 4-6 MΩ). Recordings were performed using a MultiClamp 700B amplifier interfaced with a PC through a Digidata 1440A (Molecular Devices, Sunnyvale, CA, USA). Traces were sampled at a frequency of 30 kHz and low-pass filtered at 2 kHz. Data were acquired using pClamp10 software (Molecular Devices) and analysed with Prism 9 (GraphPad Software).

### CSF analysis and treatment

After centrifugation, the supernatant and the cell pellet were stored separately at −80°C until use. Analysis of CSF levels of 69 inflammatory mediators was performed using 50 mL of supernatant CSF and a combination of immune assay multiplex techniques, based on the Luminex technology (Kit 40-Plex Human Chemokine AND Kit 37-Plex Human Inflammation Panel 1, Bio-Plex X200 System equipped with a magnetic workstation; BioRad, Hercules, CA), using Bio-Plex Software Manager 6.1. All samples were analyzed in duplicate in 2 independent experiments to verify the results’ reproducibility and consistency. The CSF level of each protein detected during the analysis was normalized to the total protein concentration of each CSF sample (measured by the Bradford protocol). For organoids exposure to CSF, SOX10-eGFP organoids stimulated with doxycycline and integrating hiPSC-derived microglia were exposed to 10% CSF for 24 hours. For RT-qPCR, ∼150 organoids from 4 different cell lines were exposed to: (1) 10% CSF from 3 healthy controls and (2) 10% CSF from 3 MS patients (**Table S6**) for 24 hours.

### 7T MRI acquisition

The *in vivo* MRI scan was obtained on a Siemens Magnetom 7T MRI scanner equipped with a birdcage-type transmit coil and a 32-channel receive coil with the following MRI sequences: (1) 2D high-resolution gradient echo sequence providing T2*-weighted and phase contrasts (TR = 1,300 ms; TE = 32 ms; 29 axial slices; FA = 50°; AT = 8 min 36 sec; in-plane resolution = 0.2 mm, slice thickness = 1 mm^3^; voxel volume = 0.04 μL); 3 minimally overlapping slabs acquired to cover most of the supratentorial brain, and (2) whole-brain 3D T1-weighted magnetization-prepared rapid gradient echo (T1-MPRAGE) (TR = 2,200 ms; TE = 2.96 ms; TI = 1,050 ms; 256 slices; FA = 7°; AT = 6 min 35 sec; nominal resolution = 0.7 mm isometric, voxel volume = 0.34 μL) before and at least 5 min after injection of 0.1 mmol/kg gadobutrol.

### Single-cell RNAseq

Details of the experiments are listed in **Table S2**. For the enzymatic cellular dissociation, 30 organoids, for each experiment, were first washed in Hank’s balanced salt solution (HBSS) and, then, incubated with a freshly prepared dissociation solution (HBSS, papain and DNAase I) for 45 min at 37°C on an orbital shaker (90 rpm), and gently triturated using a P1000 pipette. Cells were incubated again for 15 min at 37°C on an orbital shaker (90 rpm) centrifuged and gently triturated using a P1000 pipette. After the single cell dissociation, the cell suspension was mixed (1:1) with 10 mg/mL ovomucoid protease inhibitor solution and centrifuged at 300*g* for 5 min. The pellet was resuspended in 1X PBS containing BSA (1:50). Cells were counted using trypan blue and a haemocytometer under bright field. A yield of 700–1,200 cells per μL per sample was targeted. For droplet-based scRNA-seq, libraries were prepared using the Chromium Single Cell 3□ v3, according to the manufacturer’s protocol (10x Genomics) without modifications, targeting 5,000 cells per sample. Using the Illumina Novaseq platform, all samples were sequenced at 50,000 reads per cells. Preparation of cDNA libraries and sequencing support were performed at the Center for Omics Sciences (San Raffaele Scientific Hospital, Milan).

### Bioinformatic analysis

Downstream analysis utilized the computational resources of the research cluster of San Raffaele Scientific Hospital, Milan. CellRanger software (v.6.1.2; 10x Genomics) was implemented for library demultiplexing, barcode processing, fastq file generation, gene alignment to the human genome (refdata-gex-GRCh38-2020-A), and unique molecular identifier (UMI) counts. Filtered matrices were loaded in Seurat (v 4.1.1) using the Read10X function with a cut-off value of 200 unique molecular identifiers (UMIs) and min.cells parameter of 20. The filters for mitochondrial reads and for min and max total number of features was tailored for specific runs. As initial reference, the entire dataset was projected onto two-dimensional space using UMAP on the top 30 principal components. After data normalization and scaling doublets and multiplets were filtered out using DoubletFinder. After preprocessing performed on individual samples, the object was integrated using Seurat workflow (IntegrateData function). The dataset was then scaled to remove unwanted sources of variation (percent of mitochondrial reads, total counts per cells and cell-cycle score). On scaled data, linear dimension reduction (principal component analysis (PCA)) was performed, and the top 30 principal components were implemented for the unsupervised clustering. Seurat applies a graph-based approach to clustering: the FindNeighbors function implements the first 30 principal components to construct the k-nearest neighbours graph based on the Euclidean distance in PCA space, and the FindClusters function applies the Louvain algorithm to iteratively group nuclei (resolution = 0.8). The assignment of cell type identity to clusters was based on known cell lineage markers, as well as comparison with previously published and reported snRNA-seq studies. For each cluster, we extracted the average gene expression for all identified genes as well as the top 100 differentially expressed genes comparing each cluster against all the others. A differentially expressed gene was defined as a gene significantly expressed (*p* adjusted for multiple comparison, *p* < 0.05) in ≥25% of cell populations and showing, on average, >0.25-fold difference (log scale) between groups of cells.

### Donor deconvolution

The termed “no-doxy pooled sample” (experiment #1, **Table S2**) included SOX10-transfected organoids not stimulated by doxycycline from 3 distinct donors (10 organoids for each donor). To separate the contribution of the 3 donors, a donor deconvolution was performed on the single-cell RNAseq data. After sequencing, the data have been processed using CellRanger (v.6.1.2, 10x Genomics). The resulting bam file has been implemented as input of cellsnp-lite using Mode1a.^59^ cellsnp-lite is designed to pileup the expressed alleles in single-cell RNA-seq data, which can be directly used for donor deconvolution in multiplexed designs. After pileup, Vireo was used to demultiplex the donors by clustering the cells based on their single-nucleotide polymorphism (SNPs) profiles.^60^

### Cluster compositional analysis

To this end, two different algorithms were performed: MiloR and Cacoa.

1. MiloR: we performed compositional analysis on the organoids single cell transcriptomic datasets using a tool for differential abundance testing on k-nearest neighbor (KNN) graph neighborhoods, implemented in the R package miloR https://github.com/MarioniLab/miloR. In brief, we performed PCA dimensionality reduction and KNN graph embedding on the dataset. We defined a neighborhood as the group of cells connected to a sampled cell by an edge in the KNN graph. Cells were sampled for neighborhood construction and tested for differences in abundance between the cells from doxycycline-stimulated *vs* not organoids. Data are reported as density plot of the neighbors.
2. Cacoa: we performed compositional analysis on the organoids single cell transcriptomic datasets, using a tool for differential abundance testing on isometric log-ratio transformation (ilr), implemented in the R package Cacoa https://github.com/kharchenkolab/cacoa. The 17 clusters were transformed into 17-1 simplex space using isometric log-ratio transformation (ilr). As ilr transformation uses normalization by geometric mean of abundances, the resulting 16 variables were mainly independent and, thus, suitable for statistical comparisons. To compare the cluster composition in the comparison, we identified a surface within this 16-simplex space that optimally separated the two conditions and estimated the extent to which different cell types drive the orientation of that surface measured and loading coefficients. To estimate the statistical significance of loading coefficients, we generated an empirical background distribution based on random resampling of samples and cells across the entire dataset.

### Pseudotime trajectory analysis

We used Monocle to generate the pseudotime trajectory analysis using the single-cell RNAseq dataset from organoids derived from donor 3 (without hiMicroglia incorporation). This approach allows to focus only on NPC-derived cells’ differentiation trajectory. UMAP embeddings and cell subclusters generated from Seurat were converted to a CellDataSet object using SeuratWrappers and then used as input to perform trajectory graph learning and pseudotime measurement through independent component analysis (ICA) with Monocle. Heatmap of the gene expression at the branch points was produced using the function plot_genes_branched_heatmap.

### Cell-to-cell communication

To infer cell–cell communication, we used the single-cell RNAseq dataset from GIBCO and SOX10-eGFP expressing organoids (both stimulated and nonstimulated with doxycycline) incorporating hiMicroglia. Cell–cell interactions based on the expression of known ligand–receptor pairs in different cell types were inferred using CellChat (v.1.5.0).^30^ In brief, following the official workflow, we loaded the normalized counts into CellChat and applied the preprocessing functions ‘identifyOverExpressedGenes’, ‘identifyOverExpressedInteractions’ and ‘projectData’ with standard parameters set. As database we selected the ‘Secreted Signaling’ pathways and used the precompiled human ‘Protein-Protein-Interactions’ as a priori network information. For the main analyses the core functions ‘computeCommunProb’, ‘computeCommunProbPathway’ and ‘aggregateNet’ were applied using standard parameters and fixed randomization seeds. Finally, to determine the senders and receivers in the network the function ‘netAnalysis_signalingRole’ was applied on the ‘netP’ data slot.

### Reference-based mapping

To assess the transcriptional similarity between the cell types in the organoids and the CNS cells of the adult human brain, we mapped the organoids’ single cell transcriptomic data onto a reference atlas of the adult human primary motor cortex from non-neurological subjects^16,17^ using the Azimuth algorithm (https://satijalab.org/azimuth/). For each cell, a mapping score (reflecting the confidence associated with a specific annotation) and a prediction score (reflecting how well represented the cell is by the reference atlas) was generated. The mapping and the prediction scores provided by Azimuth range from 0 (no match) to 1 (perfect match). The same approach was implemented also to verify the transcriptional similarity between the cell types from 8 published single-cell RNAseq datasets^19–26^ from cortical organoids (metanalysis available in Tanaka et al.,^28^) and the Azimuth adult human primary cortex.

A third label-transfer analysis was performed using a published snRNAseq dataset from MS brain tissue^3^ to assess the transcriptional similarity between the immune cells and the astrocytes of the organoids *vs* the immune and astrocyte subclustering of the adult MS human brain. To convert the MS dataset into a reference atlas, for each subcluster of interest, the Seurat pipeline was computed by adding the “model” argument to the UMAP function.

### Gene Set Enrichment Analysis (GSEA)

We retrieved the list of differentially expressed genes of both microglia inflamed in MS (MIMS)-iron *vs* homeostatic microglia as well as MIMS-foamy *vs* homeostatic microglia (*p* adjusted < 0.05, log_2_ fold change > 0.5) from a previously published snRNAseq dataset of chronic active MS brain lesions.^3^ From the two ranked lists, we generated two signatures for the two MIMS subsets, after removing genes coding for ribosomal proteins. These signatures were then used to run Gene Set Enrichment Analysis (GSEA)^61^ on the full ranks (CSF-exposed *vs* control) for the organoids immune clusters 12 and 13.

### Differential expression and pathways analysis in MS-derived vs control-derived organoids

For each cell type, using scRNAseq pseudobulk data, we performed a differential expression analysis using Deseq2.^62^ The differential expressed genes (defined by *p* value adjusted to multiple comparison < 0.05 and abs log2 fold change >1) were used as input for the enrichment analysis using enrichR package (with Reactome and GO:BP annotations).^63^ Significant enriched terms (defined by p value adjusted to multiple comparison <0.05 and odd ratio >5) were analyzed for each cell type.

### Label-free quantitative proteomic analysis

Patient-derived organoids were lysed using M-PER™ Mammalian Protein Extraction Reagent (Thermo Scientific), added with protease (Thermo Fisher Scientific) and phosphatase inhibitor cocktails (PhoStop, Merck). In details, M-PER™ Mammalian Protein Extraction Reagent was added to a pellet of PDOs in a volume ratio 10:1. The lysates were mechanically disrupted by means of a syringe with a 25 Gauge needle. The samples were shaken at 400 rpm at 4°C for 10 minutes and then centrifuged at 14,000 rpm for 15 minutes at 4°C. The pellets were discarded, and the supernatants were subjected to BCA analysis in order to quantify the total protein content, using BSA as standard. Forty µg of total proteins from each sample were digested using the FASP (Filter Aided Sample Preparation) protocol.^64^ Briefly, proteins were diluted to 200 µL with 0.1 M Tris/HCl pH 7.4. Cysteines were reduced with 50 µL of 0.1 M DTE (1,4-dithioerythritol) in Tris/HCl pH 7.4 and incubated at 95 °C for 1 min; then samples were centrifuged on Amicon Ultra-3K at 14,000 x g for 10 min with a solution of urea 8 M. Cysteines were alkylated with 100 µl 0.05 M IAA (iodoacetamide) in urea 8 M added on the filter and laid at room temperature for 5 min; samples were centrifuged on Amicon Ultra-3K at 14,000 x g for 10 min with a solution of urea 8 M and then incubated overnight at 37 °C in the presence of trypsin 1:50 (w/w), upon dilution of urea up to 2 M with 50 mM ammonium bicarbonate buffer.

Peptide mixtures were desalted on home-made Stage Tips C18 and injected in a capillary chromatographic system (EASY-nLC™ 1000 Integrated Ultra High Pressure Nano-HPLC System, Proxeon Biosystem) for peptide separations on a 75 µm i.d. × 15 cm reverse phase silica capillary column, packed with 1.9 μm ReproSil-Pur 120 C18-AQ (Dr. Maisch GmbH, Germany). A 100 min-gradient of eluents A (pure water with 0.1% v/v formic acid) and B (acetonitrile with 0.1% v/v formic acid) was used to achieve separation (from 5% to 40% of B in 88 min, 300□nL/min flow rate). Mass Spectromer (MS) analyses were performed using a Q-Exactive mass spectrometer (Thermo Scientific, Bremen, Germany) equipped with a nano-electrospray ion source (Proxeon Biosystems). Each sample was analysed in technical triplicates. Full scan spectra were acquired with the lock-mass option, resolution set to 70,000 and mass range from m/z 300 to 2000. The ten most intense ions (charge exclusion: unassigned, 1, 6-8, >8) were selected to be fragmented (ddMS2). MS/MS spectra were acquired with resolution set to 17,500, NCE set to 25 with an isolation window of 2 m/z. All MS/MS data were analysed using Mascot (version 2.6, Matrix Science) search engine to search the human_proteome 20220525 (101,676 sequences; 41,413,969 residues). Searches were performed with the following settings: trypsin as proteolytic enzyme; 2 missed cleavages allowed; carbamidomethylation on cysteine as fixed modification; protein N-terminus-acetylation, methionine oxidation as variable modifications; mass tolerance was set to 5 ppm and to 0.02 Da for precursor and fragment ions, respectively.

All data were then analysed by MaxQuant software (v. 1.6.1.0)^65^ for label-free protein quantification based on the precursor intensity, using the above search parameters, except for ± 20 ppm for fragment ions mass tolerance. FDR of 1% was selected for both peptide and protein identification, with minimum two peptides per protein with at least one unique. The complete dataset of identified and quantified proteins was subjected to Student’s t-test in order to define significantly differently expressed proteins, with a *p* value <0.05. The statistical analysis and the subsequent hierarchical clustering were performed using an in-house developed R script. GO analyses were performed using the free-downloadable software FunRich^66^ and tissue distribution analysis was performed using David software.^67^ For enrichment analysis, we conducted GSEA using the ranks of the detected proteins. The statistical metric utilized for protein ranking was the t-statistics derived from comparing CSF-treated versus untreated organoid samples. Annotations from KEGG (REACTOME, GO:BP) were implemented to score and the enrichment calculations were performed using the fgsea package.^68^

## QUANTIFICATION AND STATISTICAL ANALYSIS

### Organoids’ immunostaining quantification

After immunolabeling, the number of cells (branched MBP^+^ OLs and HLADR^+^ microglia) was determined by counting stained cells in at least 30 organoid cryosections per experimental conditions. Counting was independently performed by two raters at the fluorescence microscope (disagreements were resolved by consensus). For the quantification of the percentage staining area (GFAP, Tuj1, Nestin, Pax6), images were analyzed with ImageJ (Fiji Version 2.0.0) in at least 10 organoid cryosections per experimental condition.

### Statistical analysis

A detailed description of the statistical analysis is provided in each corresponding figure legend. Numeric data (mean, SD, SEM, and N) were included either in the figures or in supplemental tables corresponding to the graphs. Statistical analysis used one-way analysis of variance with corresponding post-hoc tests for multiple group comparisons. For rt-qPCR analysis, gene expression values were averaged, mean-centered, and z-score-scaled and analyzed using a mixed-effects model (interaction between treatment and time). See dedicated paragraph for the details of the snRNA-seq bioinformatic analysis; top 100 differentially expressed genes in **Table S3** were defined as genes significantly expressed (*p* adjusted for multiple comparisons < 0.05) in ≥25% of cell populations, and showing, on average, >0.25-fold difference (log2 scale) between groups of cells. Analyses were performed using R 4.1.1 and Prism software (GraphPad software, San Diego, CA, USA; version 9.2.0). In all reported statistical analyses, effects were designated as significant for *p* < 0.05, and *p* values are labelled as \**p* <0.05, \*\**p* <0.01, \*\*\**p* <0.001, ****p <0.0001. Some of the graphical objects in **Figures 1 and 4** were created with BioRender.com.

## SUPPLEMENTAL TABLES

Table S1. Clinical and demographic feature of the donor of fibroblasts and year of skin biopsy, related to Figure 1.

Table S2. Details of single-cell RNAseq organoids experiments, related to Figure 2.

Table S3. Top 100 positive differentially expressed genes for each cluster against all the others, related to Figure 2.

Table S4. scRNAseq pathway analysis (Enrichr) MS-derived vs control-derived organoid cells by cell cluster (p adjusted<0.05, odds.ratio>5), related to the main text.

Table S5. Levels of 69 inflammatory mediators in the CSF implemented in the study compared by zscore to 75 CSF samples from untreated MS cases by Bio-Plex analysis, related to Figure 4.

Table S5. Pathway analysis (Enrichr) exposed *vs* non-exposed CSF assembloids by cell cluster, related to Figure 4.

Table S6. Clinical and demographic feature of the donor of cerebrospinal fluid implemented in the study, related to Figure 4.

Table S7. Bulk proteomics pathway analysis in the exposed vs non-exposed CSF organoids, related to Figure S5.

Table S8. scRNASeq pathway analysis (Enrichr) exposed vs non-exposed CSF organoids by cell cluster, related to Figure 4.

Table S9. scRNAseq pathway analysis (Enrichr) using 2 conditions: CSF-treated vs untreated and MS-derived vs control-derived organoid cells by cell cluster (p adjusted<0.05, odds.ratio>5), related to the main text.

Table S10. TaqMan assay list, related to Key Resources Table.

