## Supplementary Figures for "Glia-enriched stem-cell 3D model of the human brain mimics the glial-immune neurodegenerative phenotypes of multiple sclerosis"

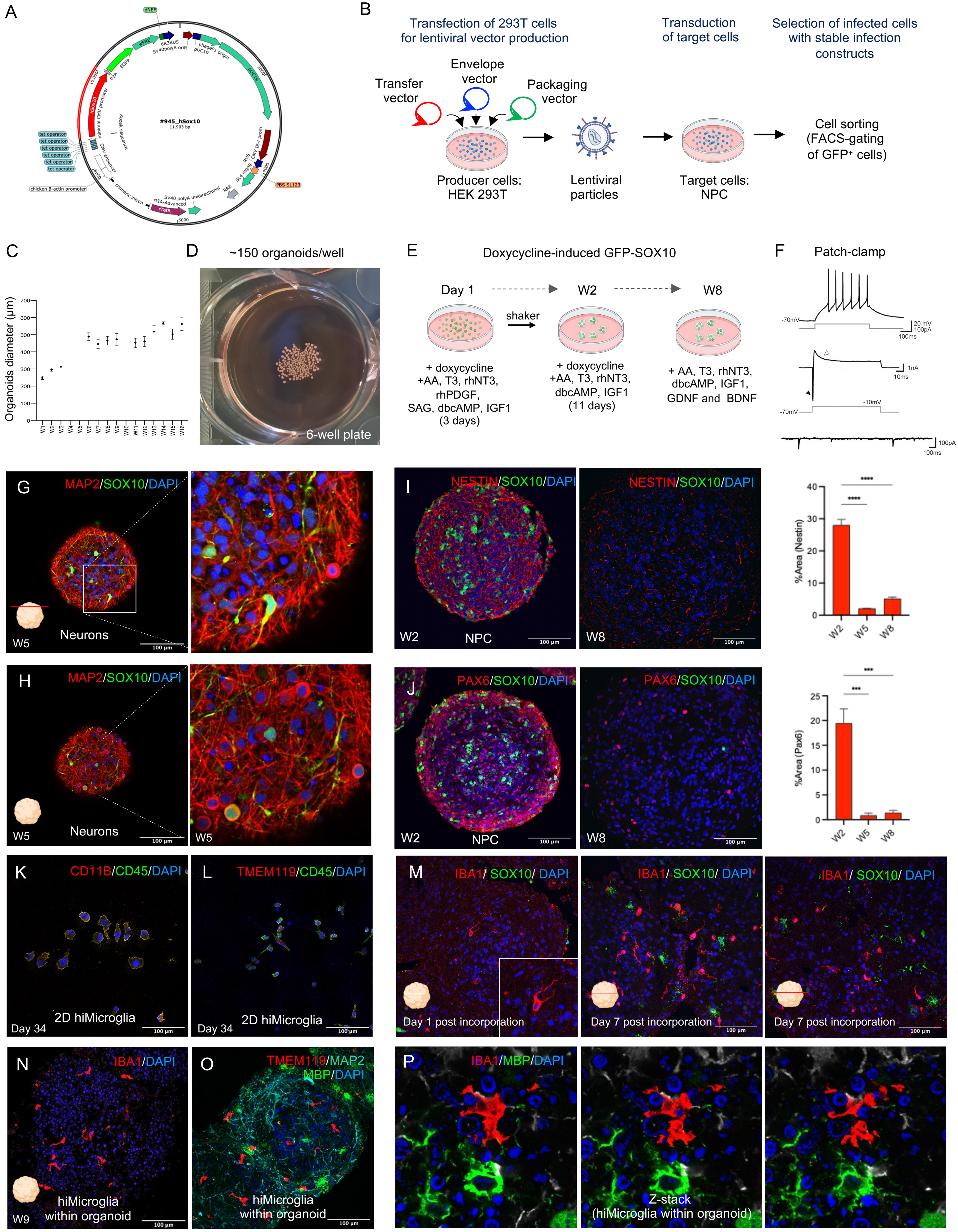
A
B
Transfection of 293T cells for lentiviral vector production
Transduction of target cells
Selection of infected cells with stable infection constructs
Transfer vector
Envelope vector
Packaging vector
Producer cells: HEK 293T
Lentiviral particles
Target cells: NPC
Cell sorting (FACS-gating of GFP+ cells)
C
Organoids diameter (µm)
D
~150 organoids/well
6-well plate
E
Doxycycline-induced GFP-SOX10
Day 1
W2
W8
shaker
+ doxycycline +AA, T3, rhNT3, rhPDGF, SAG, dbcAMP, IGF1 (3 days)
+ doxycycline +AA, T3, rhNT3, dbcAMP, IGF1 (11 days)
+ AA, T3, rhNT3, dbcAMP, IGF1, GDNF and BDNF
F
Patch-clamp
G
MAP2/SOX10/DAPI
W5
Neurons
H
MAP2/SOX10/DAPI
W5
Neurons
I
NESTIN/SOX10/DAPI
W2
NPC
W8
%Area (Nestin)
J
PAX6/SOX10/DAPI
W2
NPC
W8
%Area (Pax6)
K
CD11B/CD45/DAPI
Day 34
2D hiMicroglia
L
TMEM119/CD45/DAPI
Day 34
2D hiMicroglia
M
IBA1/ SOX10/ DAPI
Day 1 post incorporation
Day 7 post incorporation
N
IBA1/DAPI
W9
hiMicroglia within organoid
O
TMEM119/MAP2 MBP/DAPI
hiMicroglia within organoid
P
IBA1/MBP/DAPI
Z-stack (hiMicroglia within organoid)
100 µm

**Figure S1. SOX10-eGFP organoids' production and characterization by immunofluorescence. Related to Figure 1.**

(A) Lentiviral vector encoding the SOX10 coding sequence and GFP as reporter gene under the control of doxycycline-inducible promoter.

(B) Schematic for transfection of 293T HEK cells with transfer, envelope, and packaging vectors for lentiviral particles production, transduction of target cells (*i.e.,* NPCs) and selection of infected NPCs with stable infection constructs by FACS-gating of $GFP^+$ cells.

(C) Organoids' diameter (μm) over time (W=weeks) ($n$=10 organoids per time point).

(D) Representative image of organoids ($n$ ~ 150 organoids) in a well of a 6-well-plate.

(E) Schematic for doxycycline-induced SOX10 induction during the 8-week protocol (see **STAR METHODS** for a detailed description).

(F) Electrophysiological organoid recording: example of action potential firing (black trace) in response to injection of a suprathreshold current step (50 pA, 500 ms; gray trace); representative trace showing an inward $Na^+$ (black arrowhead) and an outward $K^+$ current (white arrowhead) evoked by a voltage step from -70 to -10 mV (100 ms, gray trace); example of spontaneous postsynaptic currents (excerpted from a longer trace lasting 1 minute) recorded at -60 mV in voltage-clamp mode.

(G-H) Representative images of free-floating organoids exposed to doxycycline after 5 weeks of differentiation. Immunostaining of $MAP^+$ neurons and OPCs endogenously expressing SOX10. Nuclei are marked with DAPI.

(I) Immunostaining of $NESTIN^+$ and $SOX10^+$ cells (30X) out of DAPI in whole cryosections from 2- and 8-week-old organoids exposed to doxycycline and quantification of $NESTIN^+$ cells (% staining area) at 2, 5, and 8 weeks (one-way ANOVA and multiple comparison) ($n$=3).

(J) Immunostaining of $PAX6^+$ and $SOX10^+$ cells (30X) out of DAPI in whole cryosections from 2- and 8-week-old organoids exposed to doxycycline and quantification of $PAX6^+$ cells (% area) at 2, 5, and 8 weeks ($n$=3, one-way ANOVA and Tukey's multiple comparisons test, * $p<0.05$, ** $p<0.01$, *** $p<0.001$).

(K-L) Immunolabeling of $CD11B^+$, $CD45^+$(left: 30X), $TMEM119^+$ cells (right: 30X) in 2D culture of hiMicroglia at day 34 of differentiation. Nuclei are marked with DAPI.

(M) Immunolabeling of $IBA1^+$ and $SOX10^+$ cells (30X) out of DAPI in cryosections from 8- and 9-weeks-old organoids 1 day and 7 days after hiMicroglia incorporation.

(N-O) Immunolabeling of $IBA1^+$ hiMicroglia (left: 30X) and $TMEM119^+$/$MBP^+$/$MAP2^+$ cells (right: 30X) out of DAPI in whole cryosections from 9 weeks-old organoids.

(P) Z-stacks of $IBA1^+$ microglia and $MBP^+$ OL out of DAPI in whole cryosections from 9 weeks-old organoids.

*Abbreviations:* NPCs: neural precursor cells.

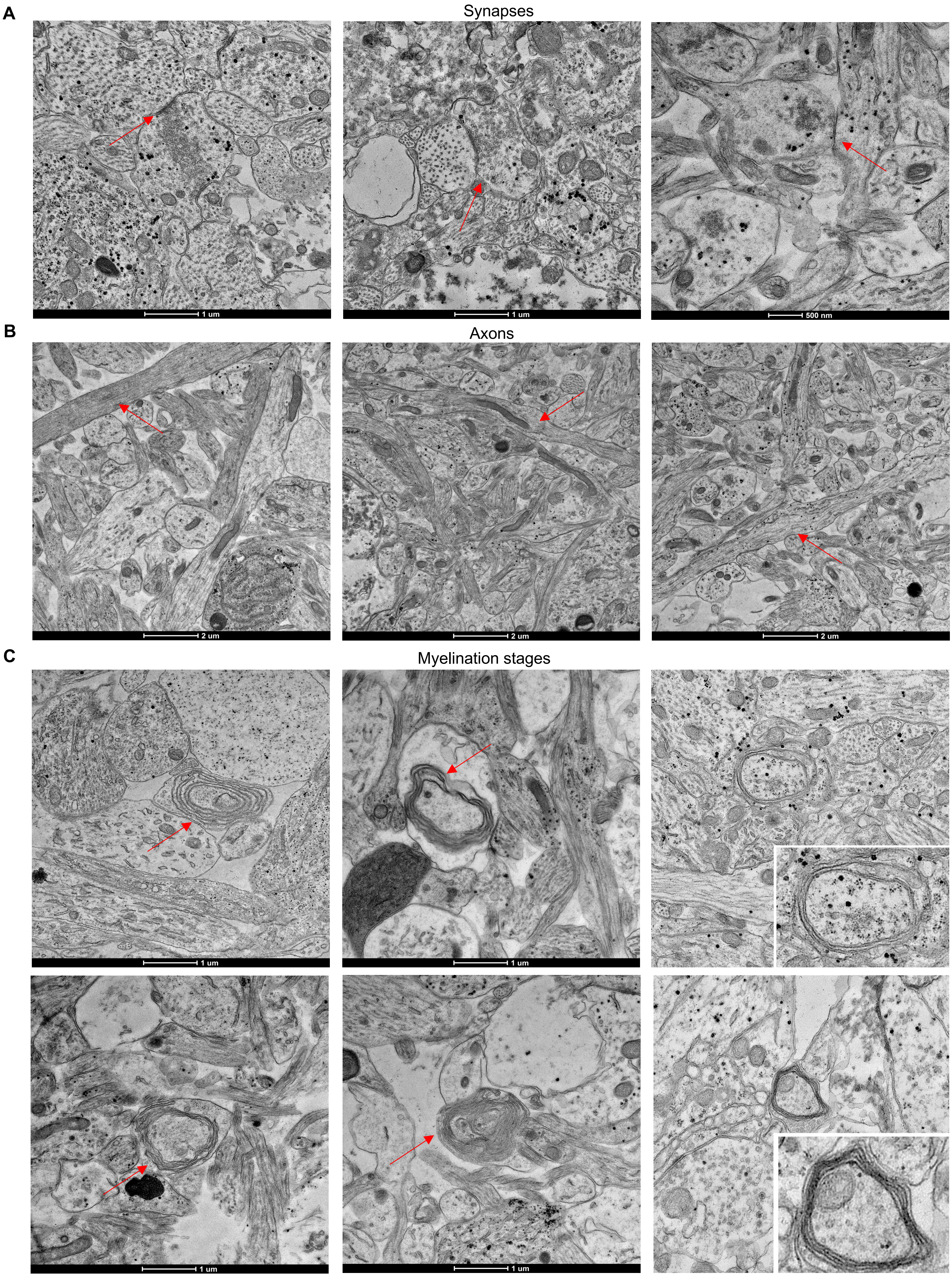
A
Synapses
1 um
1 um
500 nm
B
Axons
2 um
2 um
2 um
C
Myelination stages
1 um
1 um
1 um
1 um

**Figure S2. Ultrastructure analysis on organoids. Related to Figure 1.**
(A-B) Ultrastructure analysis by electron microscopy at 8 weeks showed: (A) evidence of cell-to-cell junctions (red arrowhead) demonstrating functional interactions between the cells in doxycycline-treated organoids; (B) axonal projections (red arrowhead) in doxycycline-treated organoids.
(C) Transmission electron microscopy images of organoids showing early myelination processes during the differentiation protocol (2 and 5 weeks) to late time points (8 weeks). Electron microscopy was performed on organoids derived from 4 independent experiments with similar results.

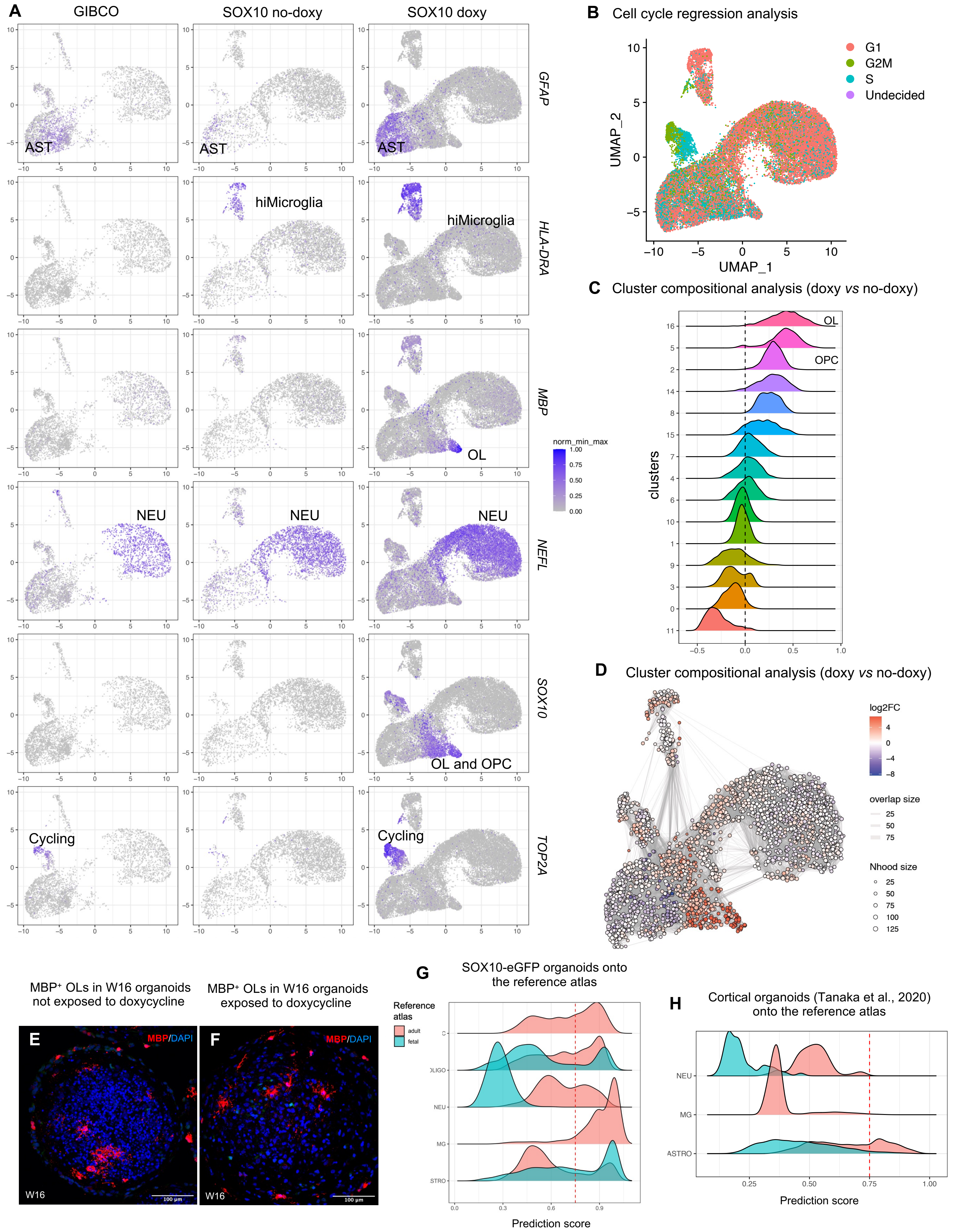

A
GIBCO
SOX10 no-doxy
SOX10 doxy
GFAP
HLA-DRA
MBP
NEFL
SOX10
TOP2A
AST
hiMicroglia
OL
NEU
OL and OPC
Cycling
norm_min_max
B
Cell cycle regression analysis
G1
G2M
S
Undecided
UMAP_1
UMAP_2
C
Cluster compositional analysis (doxy vs no-doxy)
clusters
OPC
D
Cluster compositional analysis (doxy vs no-doxy)
log2FC
overlap size
Nhood size
MBP⁺ OLs in W16 organoids not exposed to doxycycline
MBP⁺ OLs in W16 organoids exposed to doxycycline
E
F
MBP/DAPI
W16
100 µm
G
SOX10-eGFP organoids onto the reference atlas
Reference atlas
adult
fetal
NEU
MG
Prediction score
H
Cortical organoids (Tanaka et al., 2020) onto the reference atlas
ASTRO

**Figure S3. Organoids' cell type and cluster composition. Related to Figure 2.**
(A) UMAP showing *GFAP*, *HLA-DRA*, *MBP*, *NEFL, SOX10* and *TOP2A* expression in organoids.
(B) Cell cycle regression analysis indicating cells in G1, G2M, and S phase.
(C) Organoids' cluster compositional analysis for the comparison exposed *vs* not exposed to doxycycline: using Cacoa algorithm, the statistical significance of loading coefficients is reached for clusters 16, 2, 15, 7, 8, 5, and 1.
(D) Organoids' cluster compositional analysis for the comparison exposed *vs* not exposed to doxycycline: using MiloR, the density plot of the neighbors shows that the oligolineage clusters are significantly denser in doxycycline-stimulated organoids.
(E-F) Immunostaining of $MBP^+$ OL (30X) out of DAPI in whole cryosections from 16-week-old organoids not exposed to doxycycline *vs* organoids exposed to doxycycline.
(G-H) SOX10-organoids (G) and human cortical organoids (scRNAseq dataset metanalysis from Tanaka et al.) scRNAseq data were mapped on the human adult vs fetal reference atlases. The distribution of the mapping prediction scores is shown separately for adult (pink) *vs* fetal (cyan) atlas. The red dotted line indicates the prediction scores above 0.75, indicating high confidence supported by multiple consistent gene anchors.

*Abbreviations:* NEU: neurons; ASTRO: astrocytes; OPC: oligodendrocyte precursor cells; OL: oligodendrocytes; CYCLING: cycling cells.

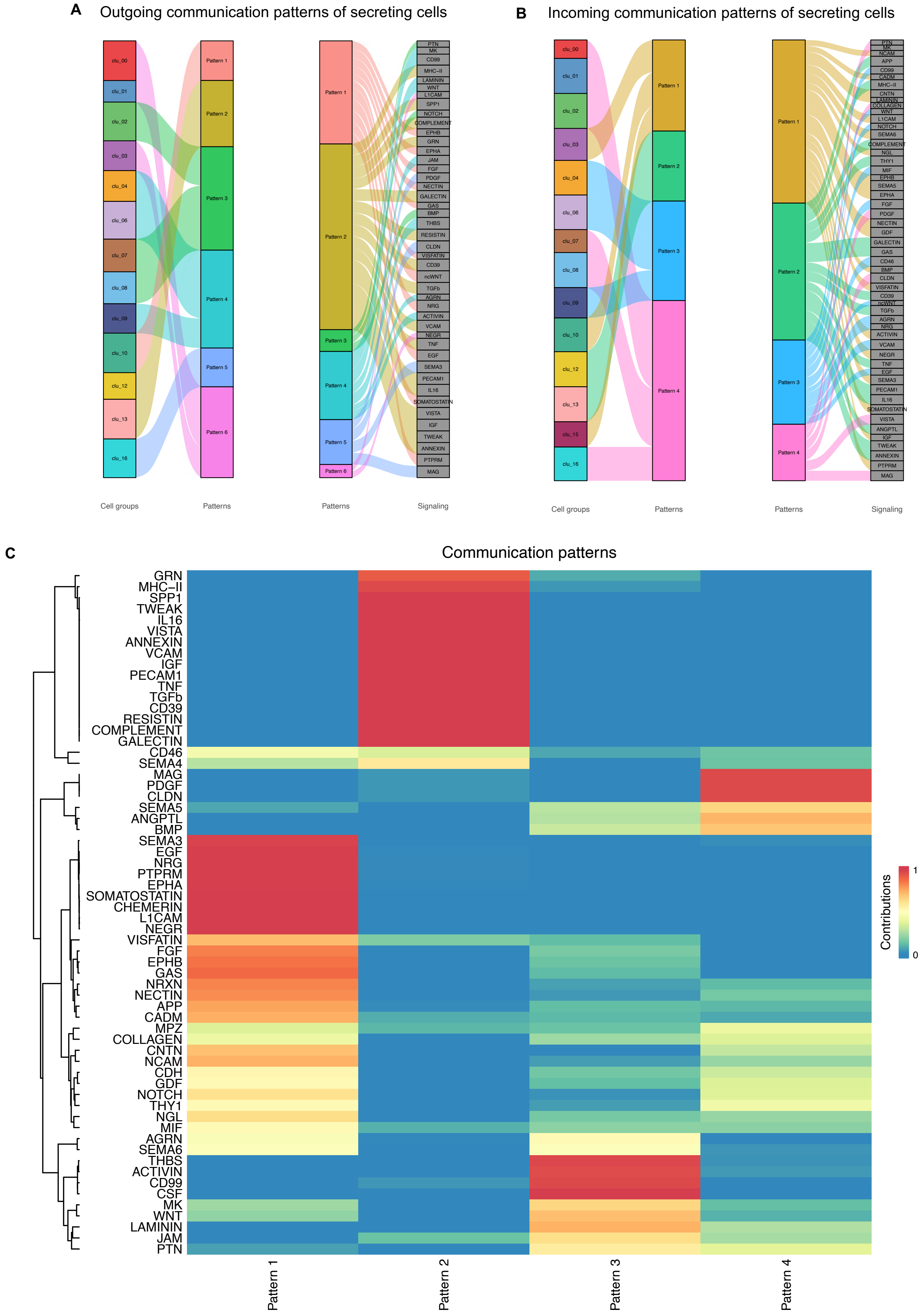
A
Outgoing communication patterns of secreting cells
clu_00
clu_01
clu_02
clu_03
clu_04
clu_06
clu_07
clu_08
clu_09
clu_10
clu_12
clu_13
clu_16
Pattern 1
Pattern 2
Pattern 3
Pattern 4
Pattern 5
Pattern 6
PTN
MK
CD99
MHC-II
LAMININ
WNT
L1CAM
SPP1
NOTCH
COMPLEMENT
EPHB
GRN
EPHA
JAM
FGF
PDGF
NECTIN
GALECTIN
GAS
BMP
THBS
RESISTIN
CLDN
VISFATIN
CD39
ncWNT
TGFb
AGRN
NRG
ACTIVIN
VCAM
NEGR
TNF
EGF
SEMA3
PECAM1
IL16
SOMATOSTATIN
VISTA
IGF
TWEAK
ANNEXIN
PTPRM
MAG
Cell groups
Patterns
Patterns
Signaling
B
Incoming communication patterns of secreting cells
clu_00
clu_01
clu_02
clu_03
clu_04
clu_06
clu_07
clu_08
clu_09
clu_10
clu_12
clu_13
clu_15
clu_16
Pattern 1
Pattern 2
Pattern 3
Pattern 4
PTN
MK
NCAM
APP
CD99
CADM
MHC-II
CNTN
LAMININ
COLLAGEN
WNT
L1CAM
NOTCH
SEMA6
COMPLEMENT
NGL
THY1
MIF
EPHB
SEMA5
EPHA
FGF
PDGF
NECTIN
GDF
GALECTIN
GAS
CD46
BMP
CLDN
VISFATIN
CD39
ncWNT
TGFb
AGRN
NRG
ACTIVIN
VCAM
NEGR
TNF
EGF
SEMA3
PECAM1
IL16
SOMATOSTATIN
VISTA
ANGPTL
IGF
TWEAK
ANNEXIN
PTPRM
MAG
Cell groups
Patterns
Patterns
Signaling
C
Communication patterns
GRN
MHC-II
SPP1
TWEAK
IL16
VISTA
ANNEXIN
VCAM
IGF
PECAM1
TNF
TGFb
CD39
RESISTIN
COMPLEMENT
GALECTIN
CD46
SEMA4
MAG
PDGF
CLDN
SEMA5
ANGPTL
BMP
SEMA3
EGF
NRG
PTPRM
EPHA
SOMATOSTATIN
CHEMERIN
L1CAM
NEGR
VISFATIN
FGF
EPHB
GAS
NRXN
NECTIN
APP
CADM
MPZ
COLLAGEN
CNTN
NCAM
CDH
GDF
NOTCH
THY1
NGL
MIF
AGRN
SEMA6
THBS
ACTIVIN
CD99
CSF
MK
WNT
LAMININ
JAM
PTN
Pattern 1
Pattern 2
Pattern 3
Pattern 4
Contributions
1
0

**Figure S4. Cell-to-cell communication in organoids. Related to Figure 3.**
(A) The inferred outgoing communication patterns of secreting cells, which shows the correspondence between the inferred latent patterns and cell groups, as well as signaling pathways. The thickness of the flow indicates the contribution of the cell group or signaling pathway to each latent pattern.
(B) The inferred incoming communication patterns of target cells that shows the correspondence between the inferred latent patterns and cell groups, as well as signaling pathways. The thickness of the flow indicates the contribution of the cell group or signaling pathway to each latent pattern.
(C) heatmap and dendrogram of the signaling pathways associated to the different cell patterns.

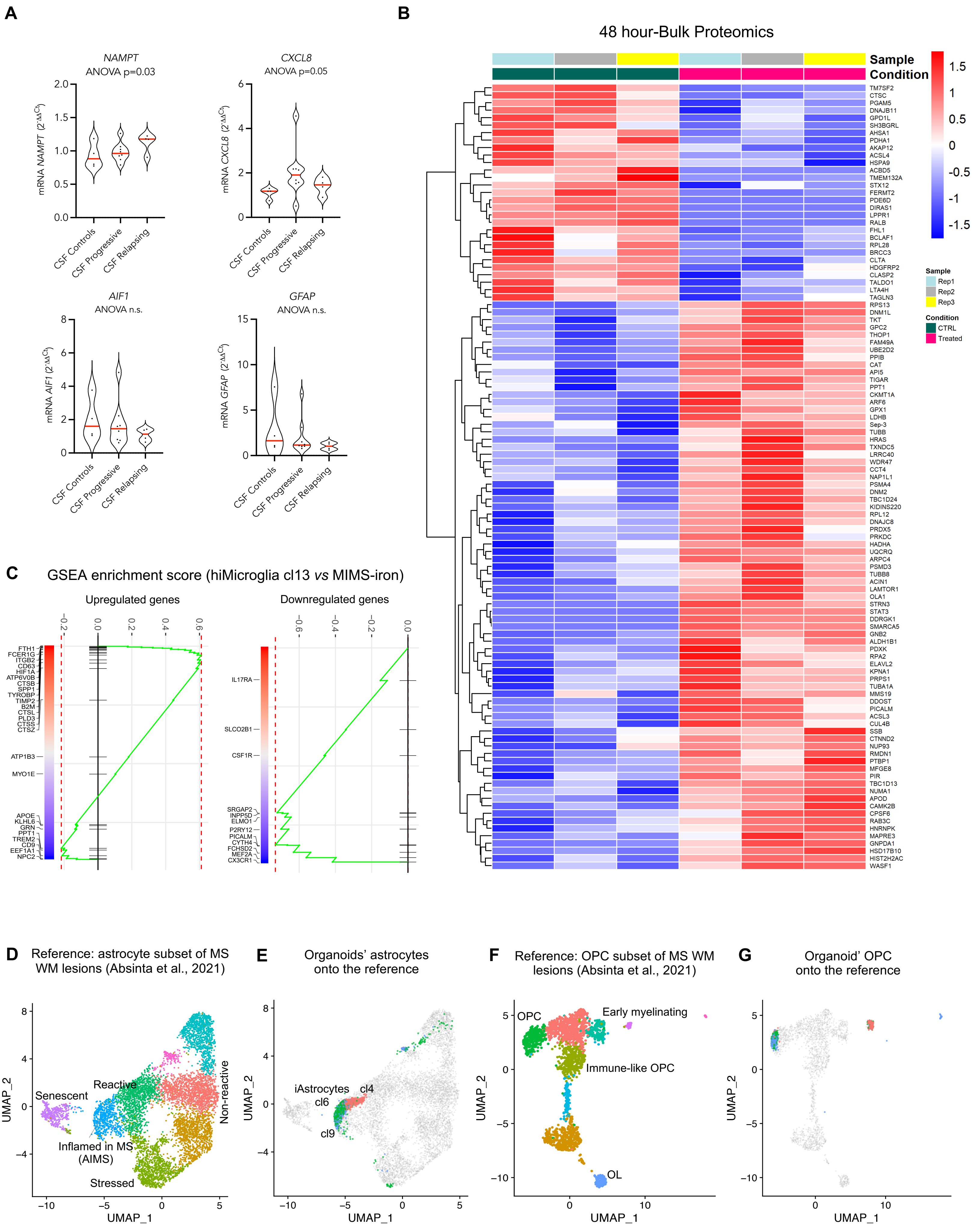

A
NAMPT
ANOVA p=0.03
mRNA NAMPT (2^-ΔΔCt)
CXCL8
ANOVA p=0.05
mRNA CXCL8 (2^-ΔΔCt)
AIF1
ANOVA n.s.
mRNA AIF1 (2^-ΔΔCt)
GFAP
ANOVA n.s.
mRNA GFAP (2^-ΔΔCt)
CSF Controls
CSF Progressive
CSF Relapsing
B
48 hour-Bulk Proteomics
Sample
Condition
Rep1
Rep2
Rep3
CTRL
Treated
TM7SF2
CTSC
PGAM5
DNAJB11
GPD1L
SH3BGRL
AHSA1
PDHA1
AKAP12
ACSL4
HSPA9
ACBD5
TMEM132A
STX12
FERMT2
PDE6D
DIRAS1
LPPR1
RALB
FHL1
BCLAF1
RPL28
BRCC3
CLTA
HDGFRP2
CLASP2
TALDO1
LTA4H
TAGLN3
RPS13
DNM1L
TKT
GPC2
THOP1
FAM49A
UBE2D2
PPIB
CAT
API5
TIGAR
PPT1
CKMT1A
ARF6
GPX1
LDHB
Sep-3
TUBB
HRAS
TXNDC5
LRRC40
WDR47
CCT4
NAP1L1
PSMA4
DNM2
TBC1D24
KIDINS220
RPL12
DNAJC8
PRDX5
PRKDC
HADHA
UQCRQ
ARPC4
PSMD3
TUBB8
ACIN1
LAMTOR1
OLA1
STRN3
STAT3
DDRGK1
SMARCA5
GNB2
ALDH1B1
PDXK
RPA2
ELAVL2
KPNA1
PRPS1
TUBA1A
MMS19
DDOST
PICALM
ACSL3
CUL4B
SSB
CTNND2
NUP93
RMDN1
PTBP1
MFGE8
PIR
TBC1D13
NUMA1
APOD
CAMK2B
CPSF6
RAB3C
HNRNPK
MAPRE3
GNPDA1
HSD17B10
HIST2H2AC
WASF1
C
GSEA enrichment score (hiMicroglia cl13 vs MIMS-iron)
Upregulated genes
Downregulated genes
FTH1
FCER1G
ITGB2
CD63
HIF1A
ATP6V0B
CTSB
SPP1
TYROBP
TIMP2
B2M
CTSL
PLD3
CTSS
CTSZ
ATP1B3
MYO1E
APOE
KLHL6
GRN
PPT1
TREM2
CD9
EEF1A1
NPC2
IL17RA
SLCO2B1
CSF1R
SRGAP2
INPP5D
ELMO1
P2RY12
PICALM
CYTH4
FCHSD2
MEF2A
CX3CR1
D
Reference: astrocyte subset of MS WM lesions (Absinta et al., 2021)
Senescent
Reactive
Non-reactive
Inflamed in MS (AIMS)
Stressed
UMAP_1
UMAP_2
E
Organoids' astrocytes onto the reference
iAstrocytes
cl6
cl4
cl9
F
Reference: OPC subset of MS WM lesions (Absinta et al., 2021)
OPC
Early myelinating
Immune-like OPC
OL
G
Organoid' OPC onto the reference

**Figure S5. Transcriptomic and proteomic of MS-CSF inflamed organoids. Related to Figure 4.**
(A) Gene expression (normalized to *GAPDH* and untreated condition) of *NAMPT, CXCL8, AIF1,* and *GFAP* in 8-week-old organoids with hiMicroglia exposed to CSF from healthy subjects (*n*=3) and subjects with diagnosed MS (*n*=3; 2 primary progressive and 1 relapsing remitting), as determined by RT-qPCR (one-way ANOVA and multiple comparison) (*n*=4 RNA samples each consisting of ~200 organoids derived from 4 hiPSC lines).
(B) Heatmap and dendrogram relative to bulk proteomic analysis conducted on untreated (named "CTRL") vs 48h CSF-treated organoids (named "Treated").
(C) Gene set enrichment analysis (GSEA) performed in hiMicroglia (cluster 13) *vs* MIMS-iron for upregulated genes left) and downregulated (right) genes. The GSEA algorithm calculates an enrichment score reflecting the degree of overrepresentation at the top or bottom of the ranked list of the genes included in a gene set in a ranked list of all genes present in our scRNA-seq datasets. A positive enrichment score indicates gene set enrichment at the top of the ranked list; a negative enrichment score indicates gene set enrichment at the bottom of the ranked list.
(D) Reference atlas based on the re-analysis of astrocyte subclustering at the chronic active MS lesion in Absinta et al.[3].
(E) Mapping of 3D organoid single cell data onto the MS astrocyte subset reference.
(F) Reference atlas was based on the re-analysis of Absinta et al.,[3] OPC subclustering at the chronic active MS lesion.
(G) Mapping of 3D organoids single cell data onto the MS OPC subset reference (F).

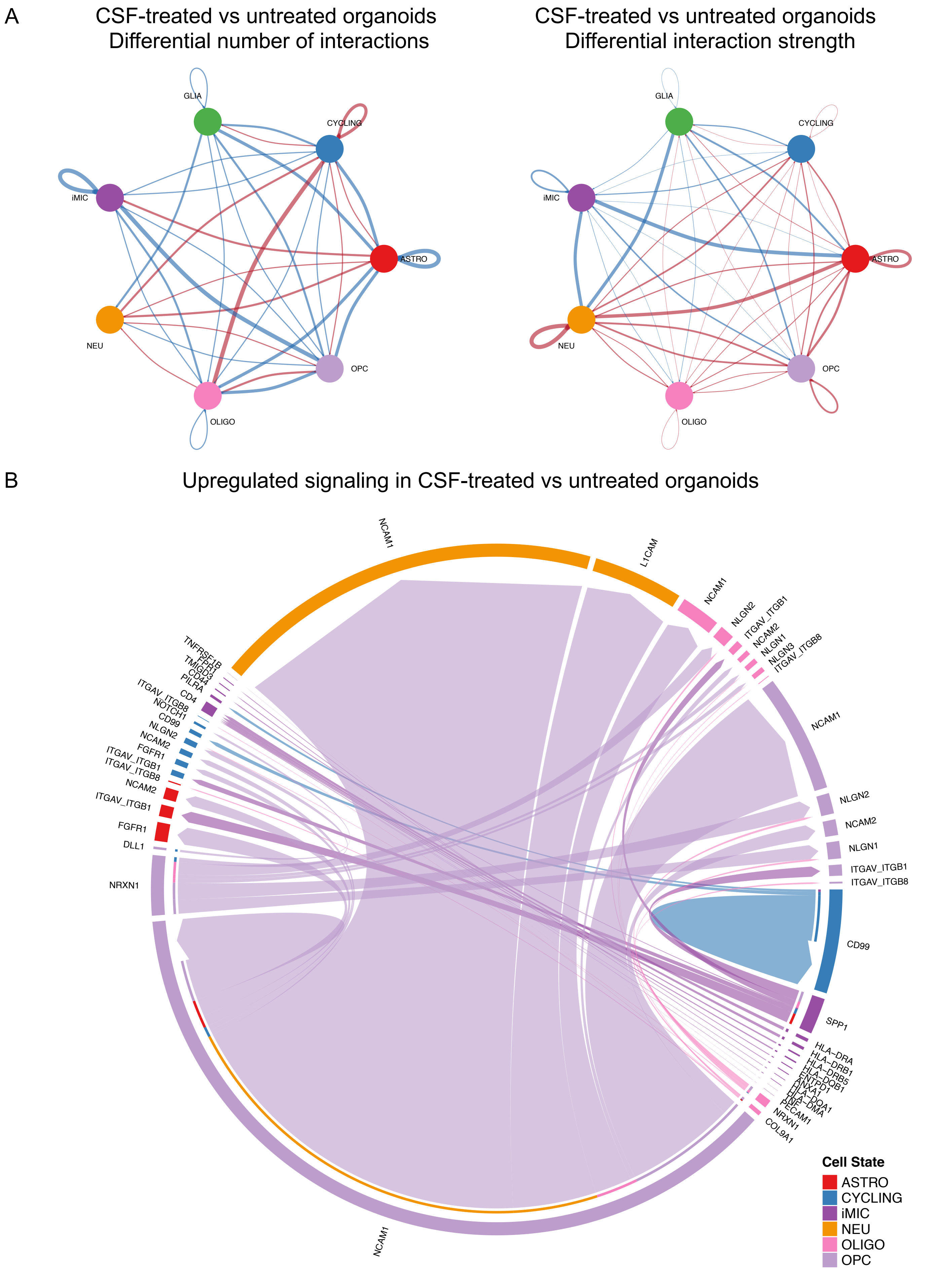
A
CSF-treated vs untreated organoids
Differential number of interactions
CSF-treated vs untreated organoids
Differential interaction strength
GLIA
CYCLING
iMIC
ASTRO
NEU
OPC
OLIGO
GLIA
CYCLING
iMIC
ASTRO
NEU
OPC
OLIGO
B
Upregulated signaling in CSF-treated vs untreated organoids
NCAM1
L1CAM
NCAM1
NLGN2
ITGAV_ITGB1
NCAM2
NLGN1
NLGN3
ITGAV_ITGB8
NCAM1
NLGN2
NCAM2
NLGN1
ITGAV_ITGB1
ITGAV_ITGB8
CD99
SPP1
HLA−DRA
HLA−DRB1
HLA−DRB5
HLA−DQB1
ENTPD1
ANXA1
HLA−DQA1
HLA−DMA
PECAM1
NRXN1
COL9A1
NCAM1
NRXN1
DLL1
FGFR1
ITGAV_ITGB1
NCAM2
ITGAV_ITGB8
ITGAV_ITGB1
FGFR1
NCAM2
NLGN2
CD99
NOTCH1
ITGAV_ITGB8
CD4
PILRA
CD44
TMIGD3
TNFRSF1B
Cell State
ASTRO
CYCLING
iMIC
NEU
OLIGO
OPC

**Figure S6. Cell-to-cell communication perturbation by MS-inflamed CSF. Related to Figure 4.**
(A) CellChat circle plots showing differential number of interactions and differential interaction strength in the crosstalk among cell populations of CSF-treated vs untreated organoids. Red lines indicate that the displayed communication is increased in CSF-treated organoids compared with untreated organoids, whereas blue lines indicate that the displayed communication is decreased in CSF-treated organoids. The line thickness is proportional to the number of interaction (circle plot on the left) or to the interaction strength (circle plot on the right).
(B) CellChat chord plot showing upregulated signaling in CSF-treated vs untreated organoids. The edge width is proportional to the indicated number of ligand-receptor pairs. Each cell population is color-coded.
